# RNA methylation influences TDP43 binding and disease pathogenesis in models of amyotrophic lateral sclerosis and frontotemporal dementia

**DOI:** 10.1101/2022.04.03.486880

**Authors:** M McMillan, N Gomez, M Bekier, X Li, R Miguez, EM Tank, SJ Barmada

## Abstract

Methylation of RNA at the N6 position of adenosine (m6A) is one of the most common RNA modifications, impacting RNA stability as well as its transport and translation. Previous studies uncovered RNA destabilization in models of amyotrophic lateral sclerosis (ALS), in association with accumulation of the RNA-binding protein TDP43, which is itself mislocalized from the nucleus in >95% of those with ALS. Here, we show that TDP43 recognizes m6A-modified RNA, and that RNA methylation is critical for both TDP43 binding and autoregulation. We also observed extensive hypermethylation of coding and non-coding transcripts in ALS spinal cord, many of which overlap with methylated TDP43 target RNAs. Emphasizing the importance of m6A for TDP43 binding and function, we identified several m6A factors that enhance or suppress TDP43-mediated toxicity via a single-cell CRISPR/Cas9 candidate-based screen in primary neurons. The most promising genetic modifier among these—the canonical m6A reader YTHDF2— accumulated within spinal motor neurons in ALS postmortem sections, and its knockdown prolonged the survival of human neurons carrying ALS-associated mutations. Collectively, these data show that m6A modifications modulate RNA binding by TDP43, and that m6A is pivotal for TDP43-related neurodegeneration in ALS.

## Introduction

Amyotrophic lateral sclerosis (ALS) is a progressive neurodegenerative disease resulting in the death of upper and lower motor neurons^1^. Limited therapeutic options exist for this condition, and the underlying pathological mechanisms remain unclear. Considerable variability in clinical, biochemical, and genetic features also complicate identification of new therapeutic targets. Despite this, over 95% of ALS cases exhibit cytoplasmic inclusions of the RNA binding protein TDP43 (TAR binding protein of 43 kDa), and mutations in *TARDBP*, the gene encoding TDP43, result in familial disease in 2-5% of individuals^2–6^. These observations imply that strategies targeting TDP43 may be relevant to the large majority of those with ALS.

TDP43 is a predominantly nuclear RNA binding protein critical for several aspects of RNA processing, including RNA splicing, transport, translation, and stability. Consistent with this, cytoplasmic mislocalization and nuclear clearing of TDP43 in ALS are closely associated with RNA missplicing, abnormal global and local mRNA translation, and widespread RNA instability^5,7–9^. We previously found that TDP43 deposition primarily destabilizes families of mRNAs encoding ribosomal proteins and oxidative phosphorylation enzymes^5^. Notably, these same mRNAs are upregulated in neurons lacking the RNA methyltransferase-like protein 14 (METTL14)^10^. This enzyme acts as a part of a “writer” complex that methylates RNA at the 6^th^ position nitrogen (N^6^-methyladenosine methylation, or m6A)^11,12^. m6A marks can be removed by demethylases (“erasers”) and/or recognized by a class of RNA binding proteins (“readers”) that, like TDP43, function in RNA splicing, transport, translation and stability^13,14^. In addition, TDP43 binds heavily methylated RNAs, and physically interacts with both writers and readers^15–18^. Together, these findings suggest that TDP43 may recognize m6A-modified RNA, raising the possibility that mislocalization and aggregation of TDP43 in ALS may preferentially affect RNA substrates carrying m6A marks.

Here, we answer the question of whether TDP43 binds m6A-modified RNA through several orthologous approaches. We show not only that the majority of TDP43 substrates carry m6A modifications, but also that the m6A reader protein YTHDF2 facilitates TDP43-related toxicity in rodent and human neuron models of ALS. Supporting the connection between TDP43 pathology in ALS and m6A RNA, we detected extensive RNA hypermethylation in postmortem spinal cord tissue from sporadic ALS patients compared to controls. These data highlight a fundamental link between m6A RNA modifications and ALS pathogenesis, potentially mediated by TDP43-dependent misprocessing of m6A-modified RNA.

## Results

### TDP43 binds m6A-modified RNA

YTH domains, common to m6A reader proteins, exhibit selective recognition of m6A-modified RNA, but previous evidence indicates that other functional domains such as RNA recognition motifs (RRMs) present in TDP43 and other members of the hnRNP family can also bind m6A-modified RNA^11,19^. Specifically, hnRNP-C^20^ and hnRNP-A2/B1^21^ are capable of recognizing m6A-modified RNA despite not having YTH domains, and the presence of m6A modifications enhances their affinity for RNA targets. Given this, we asked whether TDP43, which shows high affinity for UG rich sequences^22–24^, also recognizes m6A-modified RNA. First, we measured the UG density surrounding experimentally verified m6A sites, determined by m6A antibody cross-linking induced methylation (CIMS; **Fig. 1A**) and truncation (CITS; **Fig. 1B**)^25^. UG density is significantly greater immediately upstream of m6A sites, whereas random segments of genes of non-methylated genes showed no apparent change in UG density surrounding the m6A site **(Fig. 1A, B)**. These data indicate a possible connection between UG-rich TDP43 recognition motifs and m6A sites. To pursue this concept further and empirically determine if TDP43 recognizes m6A-modifed RNA in mammalian cells, we overexpressed TDP43 fused with HaloTag, a multifunctional adapter protein that facilitates TDP43 isolation by immunoaffinity purification^26^, in HEK293T cells. As a positive control, we also overexpressed a fusion of HaloTag and YTHDF2, a verified m6A reader protein^19^. We then isolated each protein by immunoaffinity purification, collected bound RNA and assessed total and m6A RNA by dot blot (**Fig. 1C**). TDP43-HaloTag and YTHDF2-HaloTag, but not HaloTag by itself, recognized and pulled down m6A RNA (**Fig. 1D**), showing that TDP43 and YTHDF2 are both capable of binding m6A RNA.

**Figure 1:**
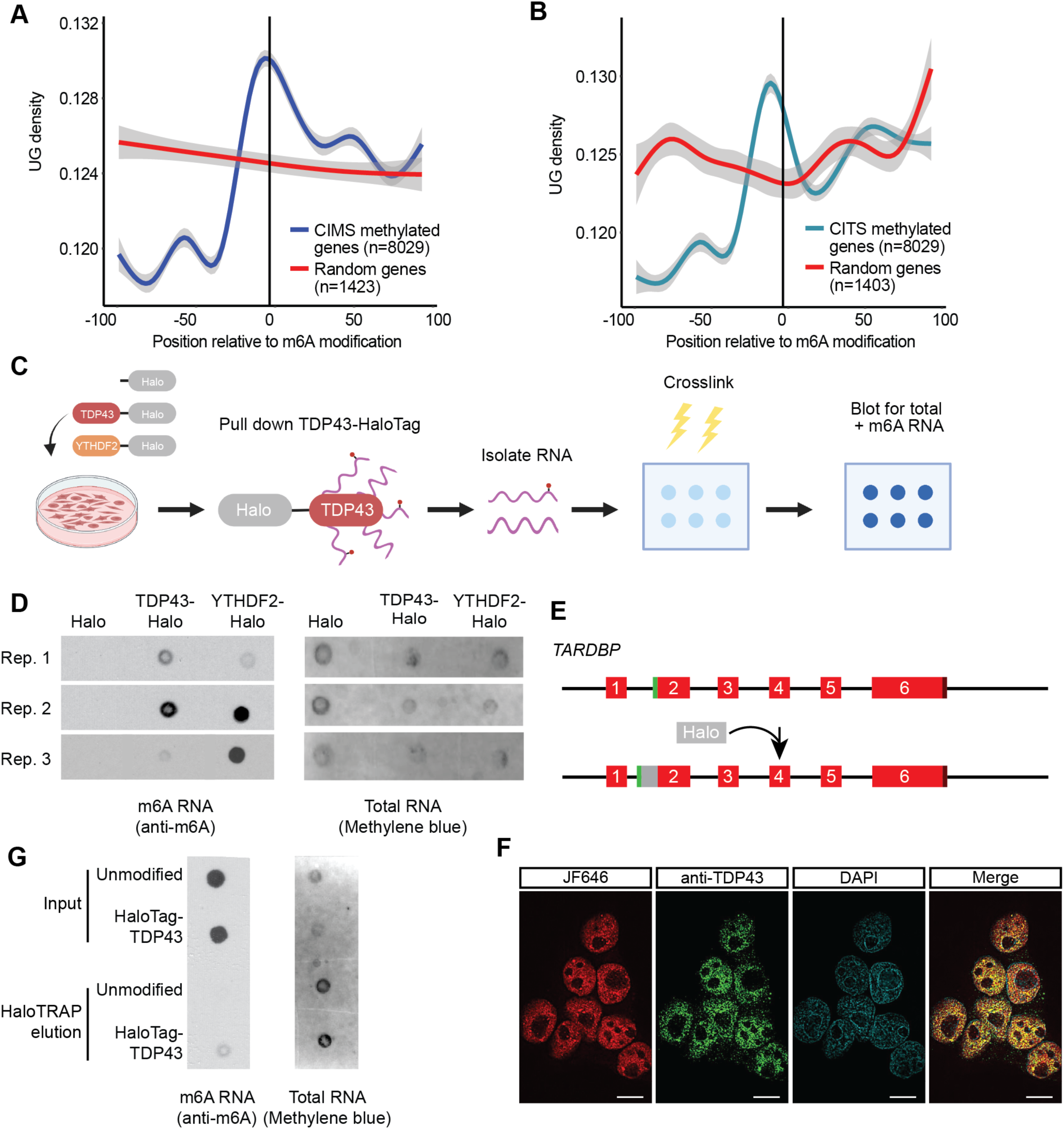
TDP43 recognizes m6A-modified RNA. Density of UG nucleotide sequences 100bp upstream and downstream of m6A modifications identified by cross-linking induced mutation sites (CIMS; **A**) or cross-linking induced truncation sites (CITS; **B**) in relation to random sequences (red line). Grey shading represents 95% confidence regions. (**C**) Schematic of HaloTag immunoprecipitation and dot blot procedure. (**D**) Dot blot for total RNA (detected by methylene blue) or m6A-modified RNA (detected by anti-m6A antibody) isolated by immunoaffinity purification of HaloTag-labeled proteins in HEK293T cells overexpressing HaloTag, TDP43-HaloTag or YTHDF2-HaloTag from 3 biological replicates. (**E**) Diagram illustrating insertion of the HaloTag open reading frame into the endogenous *TARDBP* locus immediately 5’ to the TDP43 start codon, resulting in a fusion of HaloTag to the N-terminus of TDP43. (**F**) Halo-TDP43 HEK293T cells labeled live with JF646 Halo dye (red), then fixed, permeabilized, and immunostained with anti-TDP43 antibody (green) prior to imaging. DAPI (blue) marks the nucleus of each cell. Scale bar = 10µm. (**G**) Dot blot for total RNA (detected by methylene blue) or m6A-modified RNA (detected by anti-m6A antibody) isolated by immunoaffinity purification of endogenous HaloTag-TDP43 or exogenous HaloTag. Additional replicates shown in Sup. Fig. 1.

Because protein overexpression can introduce non-specific and potentially non-physiological interactions, we engineered a line of HEK293T cells in which endogenous TDP43 was labeled at the N-terminus with HaloTag (**Fig. 1E**). Correct insertion of HaloTag into the *TARDBP* locus was verified by Sanger sequencing, and analysis of HaloTag-TDP43 function using the *CFTR* splicing reporter uncovered no loss-of-function effects associated with the fusion (**Sup. Fig. 1A**). In addition, live-cell labeling of HaloTag-TDP43 HEK293T cells with JF646, a farred cell-permeable dye that binds covalently to HaloTag, showed a strong overlap with TDP43 localization, as judged by immunostaining using TDP43 antibodies (**Fig. 1F**). We then utilized these cells to determine if endogenous HaloTag-TDP43 is capable of recognizing m6A modified RNA. As before, we detected m6A modified RNA via dot blot among transcripts pulled down by HaloTag-TDP43 (**Fig. 1G and Sup. Fig. 1B-C**), confirming that endogenously expressed TDP43 is capable of binding m6A modified RNA.

### TDP43 substrates are enriched in m6A modifications

To explicitly define the identity, number, and location of m6A modifications in TDP43 substrates, we took advantage of DART-seq (<u>d</u>eamination <u>a</u>djacent to <u>R</u>NA modification <u>t</u>argets, followed by next generation RNA <u>seq</u>uencing)^27^ (**Fig. 2A**). This method highlights specific m6A modifications without the need for m6A antibodies, which may have limited sensitivity and specificity in discriminating m6A from other RNA modifications (i.e., m6Am). Briefly, DART-seq involves the overexpression of a chimeric fusion protein consisting of the m6A-binding YTH domain, and the deaminating enzyme APOBEC1. As illustrated previously^11,12,25,27^, m6A modifications occur in the context of a highly conserved motif (DRACH, where D can be A, G or T; R is A or G; and H is A, C or T). Within this motif, the methylated adenosine (A) residue is followed by an obligate cytosine (C). Recognition of m6A by the APOBEC1-YTH fusion results in deamination of the C immediately 3’ to the modified A residue, creating a uracil (U) base that is read as a thymine (T) during sequencing. Thus, C-T transitions occurring within a DRACH motif immediately adjacent to an A signify m6A modifications. In combination with immunoaffinity purification of endogenous HaloTag-TDP43 from the HEK293T cells we created, DART-seq enables us to define not only the RNA substrates of native HaloTag-TDP43, but also the location and m6A modification status of each substrate.

**Figure 2:**
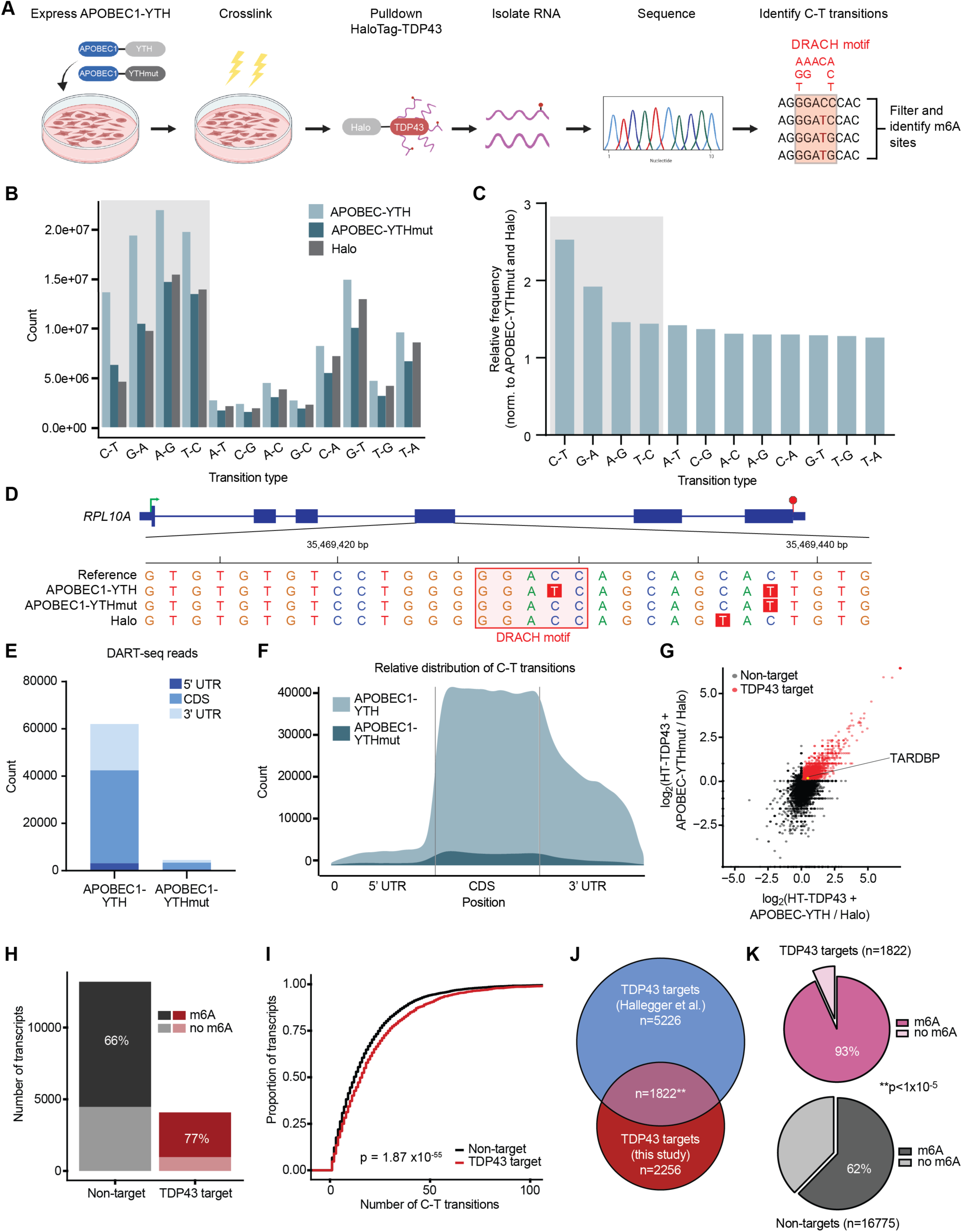
Site-specific identification of m6A-modified TDP43 substrates. (**A**) HaloTag-TDP43 immunoprecipitation was followed by DART-seq to delineate m6A sites within TDP43 target RNAs. HaloTag-TDP43 HEK293T cells were transfected with APOBEC1-YTH or APOBEC1-YTHmut and crosslinked before immunoaffinity purification of HaloTag-labeled proteins. Immunoprecipitated RNAs were then sequenced and C-T transitions were identified in the context of DRACH motifs (red shaded box, D=A/G/T, R=A/G, H=A/C/T). Absolute counts (**B**) and relative frequency (**C**) of base pair transitions observed by RNA-seq in each condition. Shaded boxes represent transition types expected from APOBEC1 activity. (**D**) Example m6A sites identified by DART-seq in *RPL10A*. C-T transitions are highlighted in red, and DRACH motifs in pink. Green arrow, transcription start site; red hexagon, transcription stop site; thick blue bars, coding exons; thin blue bars, untranslated region. (**E**) Absolute count and relative distribution (**F**) of DART-seq reads in cells expressing APOBEC1-YTH and APOBEC1-YTHmut. UTR, untranslated region; CDS, coding sequence. (**G**) Scatter plot of TDP43 targets, determined by fold enrichment in precipitated RNA from HaloTag-TDP43 cells (expressing APOBEC1-YTH and APOBEC10YTHmut) compared to cells transfected with HaloTag. Red dots signify transcripts showing ≥2-fold enrichment in both APOBEC1-YTH and APOBEC1-YTHmut expressing cells. *TARDBP*, yellow dot, identified as high confidence target. (**H**) Stacked bar graph showing percentage of m6A modified RNA in TDP43 targets (red) and non-targets (black). (**I**) Cumulative distribution of RNA methylation in TDP43 targets (red) and non-targets (black). p = 1.87×10^−55^ by Kolmogorov Smirnov test. (**J**) Euler diagram depicting overlap between TDP43 targets identified in this study, and those identified by TDP43 cross linking and immunoprecipitation followed by RNA-sequencing (CLIP-seq) in HEK293T cells (Hallegger *et al*., 2021)^28^. **p=1.5×10^−117^, hypergeometric test. (**K**) Pie charts demonstrating the percentage of methylated RNA among TDP43 targets (pink) and non-targets (grey). **p<1×10^−5^ chi-square test.

For these experiments, we transfected HaloTag-TDP43 HEK293T cells with APOBEC1-YTH or a mutated variant of APOBEC1-YTH that is unable to bind m6A modified RNA (APOBEC1-YTHmut). To control for non-specific RNA binding by HaloTag, we also transfected separate cultures of unmodified HEK293T cells with HaloTag alone. We then isolated HaloTag and HaloTag-TDP43 by immunoaffinity purification, sequenced the RNA that was pulled down in each case, and compared the resulting data to a reference database to identify base-pair transitions (**Fig. 2A**). As expected based on the deaminase activity of APOBEC1, C-T and G-A transitions— and to a lesser extent antisense T-C and A-G transitions—were strongly enriched in APOBEC1-YTH expressing cells in comparison to those expressing APOBEC1-YTHmut or HaloTag alone (**Fig. 2B, C**). These transitions occurred both within and outside of canonical DRACH motifs (**Fig. 2D**), but to increase specificity and confidence in the detection of verified m6A sites, we limited further analyses to transitions that arose in the context of a DRACH motif. In doing so, we noted profound enrichment for high-confidence m6A sites in cells expressing APOBEC1-YTH, compared to those expressing APOBEC1-YTHmut and HaloTag (**Fig. 2E, F**). The majority of detected m6A sites fall within the coding sequence (CDS) of HaloTag-TDP43 target RNAs, rather than the untranslated regions (5’ or 3’UTRs) or intronic segments (**Sup. Fig. 2A, B**). This pattern is consistent with the previously observed exclusion of m6A sites from introns, but differs from the expected concentration of m6A modifications in the 3’UTR immediately downstream of the stop codon^12,25,27^. We suspect that the unusual distribution of m6A sites likely reflects the differences between HaloTag-TDP43 substrates investigated here, and total mRNA used in prior studies.

Emphasizing the sensitivity of this approach for capturing TDP43 target RNAs, we identified 2,256 transcripts that were significantly enriched by HaloTag-TDP43 pulldown in APOBEC1-YTH and APOBEC1-YTHmut expressing cells, compared to cells expressing HaloTag alone (**Fig. 2G**). In keeping with our previous data suggesting that TDP43 recognizes m6A-modified RNA, the majority of HaloTag-TDP43 target RNAs were methylated (**Fig. 2H**), and also displayed a higher degree of methylation in comparison to non-targets. (**Fig. 2I**). To ensure that our results are not confounded by potential non-specific RNA-protein interactions of HaloTag or the immunoaffinity purification itself, we compared our HaloTag-TDP43 substrates to a published dataset of TDP43 targets in HEK293 cells identified by CLIP-seq^28^. In the process, we identified 1822 high-confidence TDP43 substrates (p= 1.5×10^−117^ for the overlap, hypergeometric test), over 90% (1699) of which were m6A-modified (**Fig. 2J, K**). By gene ontology (GO), methylated TDP43 target RNAs were highly enriched for components of the nuclear pore complex (**Sup. Fig. 3A**), a structure with intricate ties to ALS pathogenesis. Furthermore, protein-protein interaction (PPI) network prediction using STRING^29^ indicated strong enrichment for several additional disease-associated pathways, including apoptosis and p53 signaling^30^, the ribosome^5^, long-term potentiation^31^, VEGF signaling^32^ and RNA transport^33,34^ (**Sup. Fig. 3B-F**). In comparison, only 62% of non-targets were methylated (p < 1×10^−5^ for the comparison with TDP43 targets; chi-square test). These results not only confirm that TDP43 recognizes m6A modified RNA, but also indicate a possible preference of TDP43 for methylated transcripts.

### RNA methylation modulates TDP43 binding and autoregulation

To explore the functional impact of RNA methylation on TDP43 binding and function, we re-examined the DART-seq results, concentrating on the relationship between identified m6A sites and UG-rich TDP43 recognition motifs. In doing so, we observed a striking overlap between m6A modifications and UG-rich sequences primarily within the 3’UTR of HaloTag-TDP43 substrates (**Sup. Fig. 2C**); indeed, 74% of m6A sites located within 20 nt of a TDP43 motif are found within the 3’UTR of HaloTag-TDP43 targets (**Sup. Fig. 2D**). This subset of 237 transcripts was highly enriched for genes whose expression is regulated by TDP43 (**Sup. Fig. 2E**), emphasizing the functional relevance of m6A sites for TDP43-dependent regulation.

We next focused on *TARDBP*, a HaloTag-TDP43 substrate RNA that showed prominent methylation based on our DART-seq experiments. The *TARDBP* transcript, which encodes TDP43 itself, exhibited a clear C-T transition within the context of a 3’UTR DRACH motif (**Fig. 3A**). We focused on this m6A site for three reasons: First, it is located within the TDP43 binding region (TBR)^35^, a section of the 3’UTR that is both necessary and sufficient for TDP43 recognition. Second, the m6A site is immediately adjacent to a UG-rich stretch resembling the consensus GUGUGU motif commonly recognized by TDP43^36,37^. Third, the m6A site lies within a 34 nucleotide stretch (denoted CLIP34nt) that binds TDP43 *in vitro* and *in cellulo*^38^ (**Fig. 3A**).

**Figure 3:**
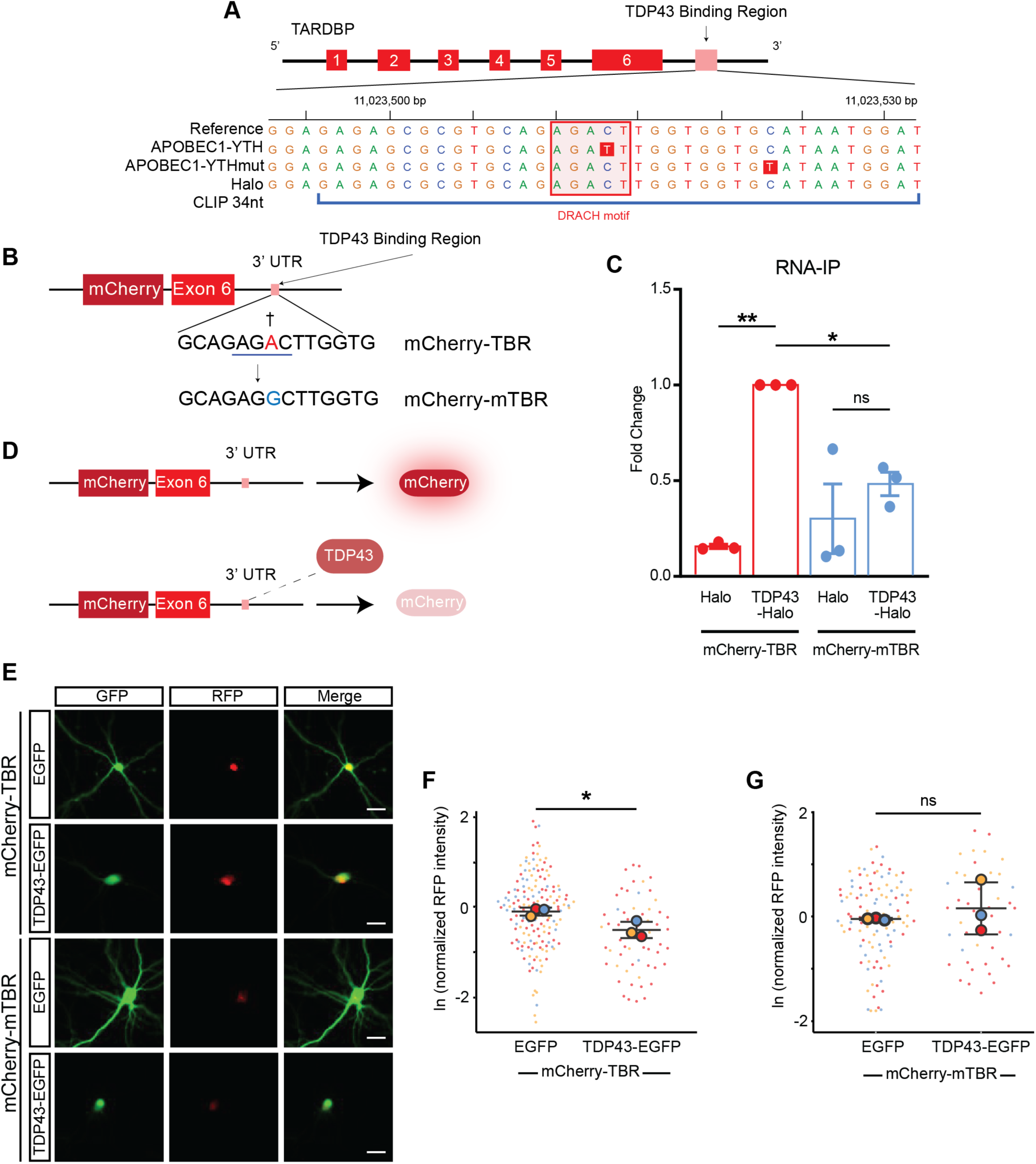
m6A modifications influence TDP43 binding and autoregulation. (**A**) *TARDBP* gene map, illustrating TDP43 binding region (TBR), the location of the DRACH motif (pink square), and the C-T transition (red box) identified by DART-seq within this domain, representing an m6A site. (**B**) Schematic of the *TARDBP* minigene reporter, consisting of the mCherry ORF upstream of *TARDBP* exon 6 and 3.4 Kb of the *TARDBP* 3’ UTR. The A residue adjacent to the detected C-T transition via DART-seq in the WT reporter (mCherry-TBR) was mutated to a G, precluding methylation the mutant reporter (mCherry-mTBR). Red, methylated residue; blue line, DRACH motif; dagger, C-T transition from DART-seq. (**C**) HaloTag-TDP43 was isolated by immunoaffinity purification from HaloTag-TDP43 HEK293T cells expressing mCherry-TBR or mCherry-mTBR, and reporter RNA detected in elution fractions by qRT-PCR. (**D**) Outline of TDP43 autoregulation assay. Excess TDP43 binds to the reporter, triggering reporter splicing, destabilization, and reduced mCherry fluorescence. (**E**) Primary rodent neurons were transfected with WT (mCherry-TBR) or mutant (mCherry-mTBR) reporters, together with EGFP or TDP43-EGFP. After 7d, mCherry expression was assessed by fluorescence microscopy. Scale bar= 20 µm. Normalized RFP (mCherry) intensity in primary neurons expressing WT mCherry-TBR reporter (**F**) or mutant mCherry-mTBR (**G**) reporter together with EGFP or TDP43(WT)-EGFP. Cherry-TBR+GFP n= 160, Cherry-TBR+TDP43(WT)-GFP n= 58, Cherry-mTBR+GFP n= 105, Cherry-mTBR+TDP43(WT)-GFP n= 44. Data in **C** plotted as mean ± SD, collected from 3 biological replicates. ns= not significant, *p< 0.05, **p< 0.01; one-way ANOVA with Tukey’s test. Data in **F** and **G** plotted as mean ± SD, color coded by biological replicate. ns = not significant, *p < 0.05; Welch’s t-test.

To probe the importance of this m6A modification for TDP43 binding and function, we utilized a minigene reporter in which *TARDBP* exon 6 and a portion of the 3’UTR containing the TBR is fused to mCherry (**Fig. 3B**; mCherry-TBR)^7,39,40^. We then applied site-directed mutagenesis to change the methylated adenosine residue shown in **Fig. 3A** to a guanosine, thereby blocking methylation of the mutated reporter (mCherry-mTBR). Both reporters were then expressed in HaloTag-TDP43 HEK293T cells or HEK293T cells transfected with HaloTag alone. Endogenous HaloTag-TDP43 was isolated by immunoaffinity purification, and bound RNA separated and assessed by qRT-PCR (**Fig. 3C**). HaloTag-TDP43 efficiently pulled down the mCherry-TBR reporter but not the m6A-deficient mCherry-mTBR reporter despite no change in reporter input levels (**Sup. Fig. 4A**), demonstrating that the m6A modification is required for recognition by TDP43.

Two possibilities may account for the apparent effect of m6A modifications on RNA recognition by TDP43: a direct influence on binding, resulting from an increase in affinity of TDP43 for m6A *vs*. unmodified RNA; or an indirect effect arising from enhanced accessibility of the TDP43 binding motif in m6A-modified RNA. Several observations argue for an indirect rather than a direct effect of m6A modifications on TDP43 binding: First, TDP43 typically recognizes UG-rich motifs rather than the DRACH sequence associated with m6A^36,37^. Second, TDP43 exhibits canonical RRM motifs rather than the more m6A-specific YTH domains found in most direct m6A reader proteins^41^. Third, hnRNP-C^20^, a heterogeneous ribonucleoprotein that is structurally and functionally related to TDP43, is an indirect m6A binding protein. To examine these possibilities in more detail, we created short (14 nt) synthetic RNA probes corresponding to the TBR with or without methylation of the predicted m6A site and performed electromobility shift assays (EMSAs) with recombinant TDP43, enabling measurements of probe binding *in vitro* (**Sup. Fig. 4B, C**). In contrast to results from RNA-immunoprecipitation studies (**Fig. 3B**), methylation of the TBR probe had little effect on TDP43 binding. In fact, we observed a slight (∼2-fold) reduction in binding affinity for the m6A-modified probe in comparison to the unmodified probe. This discrepancy could arise from differences in secondary structure between RNA transcripts present in cells and the short 14 nt probe used in these experiments, or the absence of key cellular components absent from the *in vitro* assay. Regardless, these studies suggest that m6A modifications indirectly affect TDP43 binding, perhaps by enabling greater access to UG-rich motifs buried within the TBR.

One consequence of TDP43 binding to the TBR is downregulation of TDP43 expression— this autoregulatory feedback loop is crucial for maintaining TDP43 homeostasis^6,36,42–45^. The mCherry-TBR reporter includes the majority of exon 6 and the *TARDBP* 3’UTR, facilitating investigations of TDP43 autoregulation in response to engineered mutations as well as genetic modulators of TDP43 function and toxicity^7,39,40^. Binding of TDP43 to the TBR in the mCherry-TBR reporter results in 3’UTR splicing and a reduction of mCherry fluorescence (**Fig. 3D**), likely due to a combination of nuclear retention of the spliced transcript and nonsense mediated mRNA decay (NMD)^44^. Consistent with this, TDP43-EGFP overexpression reduces mCherry fluorescence in primary neurons transfected with the mCherry-TBR reporter (**Fig. 3E, F; Sup. Fig. 4D**). In contrast, the fluorescence intensity of the m6A-deficient mCherry-mTBR reporter was unaffected by TDP43-EGFP expression (**Fig. 3G; Sup. Fig. 4D**). Thus, not only is the TBR m6A site required for recognition of the mCherry-TBR reporter, but it also affects the efficiency of TDP43 autoregulation.

### RNA hypermethylation in ALS spinal cord

Based on data in **Figs. 1** and **2** suggesting that TDP43 binds methylated RNA, and the prominent effect of RNA methylation on TDP43 autoregulation (**Fig. 3**), we questioned whether RNA methylation may be disrupted in sporadic ALS (sALS), a disorder characterized by nuclear clearance and cytoplasmic accumulation of TDP43^2–4^. To answer this question, we obtained fresh frozen spinal cord from 4 sALS patients and 3 age-matched controls (**Table S1**), isolated RNA from these samples, and quantitatively assessed transcript methylation using an m6A array (**Fig. 4A**).

**Figure 4:**
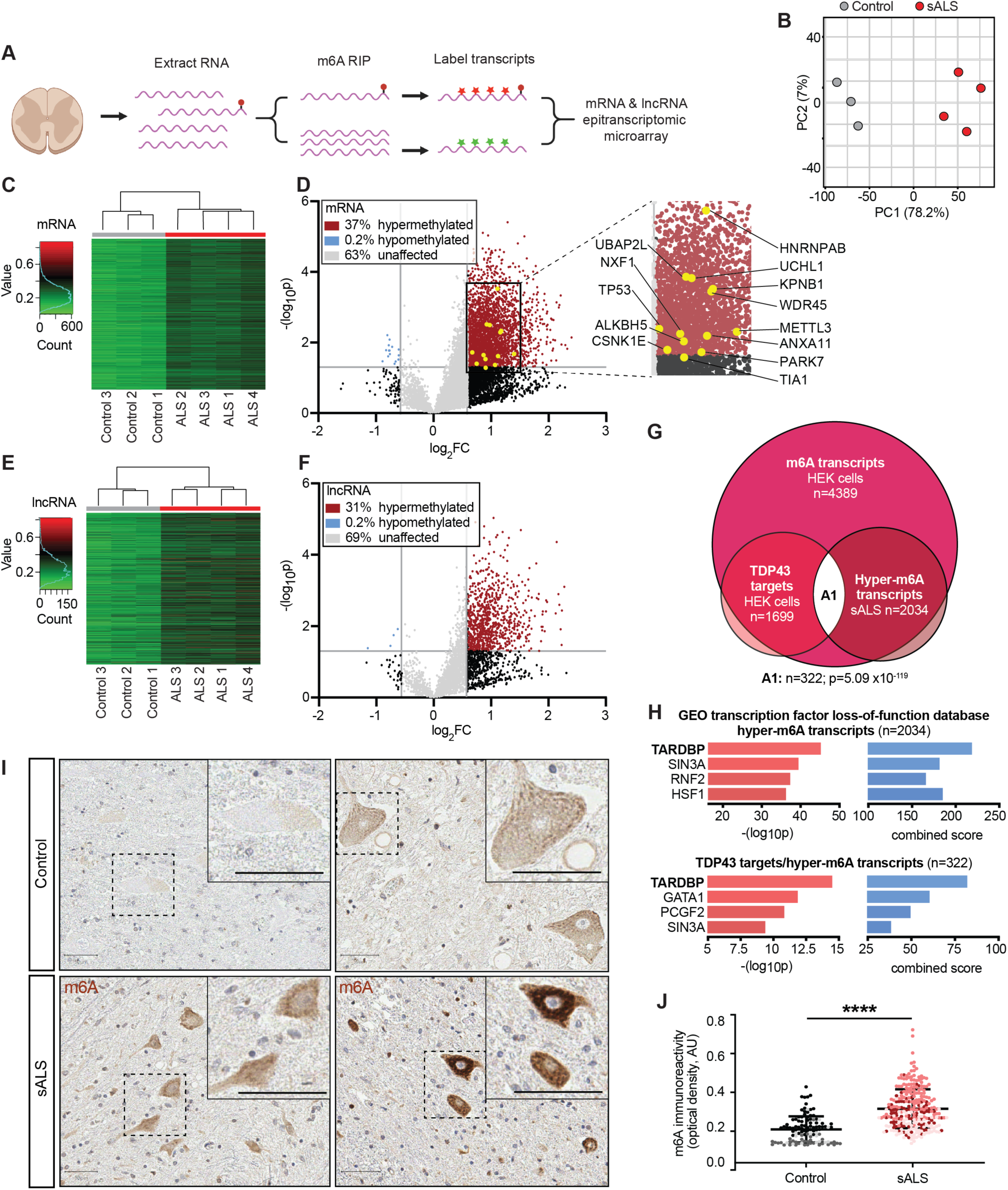
RNA hypermethylation in ALS patient spinal cord. (**A**) Genome-wide analysis of RNA methylation via epitranscriptomic array. RNA was extracted from control (n= 3) and sporadic ALS (sALS) patient (n= 4) spinal cord samples, prior to m6A RNA immunoprecipitation. The resulting samples were separated into methylated and non-methylated RNA, then labeled with distinct fluorescent dyes (red and green stars) prior to hybridization, allowing relative quantification of methylation at each annotated locus. (**B**) Principal component analysis (PCA) plot comparing methylation levels of control (grey) and ALS (red) patient samples. (**C**) Hierarchical clustering of mRNA methylation profiles from control and ALS mRNA samples. (**D**) Volcano plot depicting fold change in mRNA methylation levels in ALS compared to control spinal cord. (**E**) Hierarchical clustering of lncRNA methylation profiles from control ALS lncRNA samples. (**F**) Volcano plot showing fold change in lncRNA methylation levels in ALS compared to control spinal cord. In **D** and **F**, grey horizontal vertical lines represent p= 0.05 and fold change (FC)= 2. (**G**) Euler diagram demonstrating overlap (n= 322, p= 5.09×10^−119^, hypergeometric test) among TDP43 substrates and methylated transcripts identified in HEK293T cells, in additional to hypermethylated transcripts determined via m6A array in sALS spinal cord. Comparisons were limited to the subset of transcripts expressed in both HEK293T cells and human spinal cord (nTPM>2). (**H**) Based on comparisons with the GEO transcription factor loss-of-function database via Enrichr^83^, there was strong enrichment for TDP43-regulated genes not only among the set of 2034 transcripts hypermethylated in sALS spinal cord, but also among the 322 TDP43 targets that were also hypermethylated in sALS (A1 in **G**). Combined score = (log_10_p * Z-score). (**I**) Immunohistochemical staining for m6A in control and sALS spinal cord sections. Scale bars= 50 µm. (**J**) Quantification of m6A antibody reactivity in spinal cord neurons from control (n= 110 neurons) and sALS (n= 277 neurons) sections. Plot shows mean +/- SD, color coded by patient. ****p< 0.0001 via Mann-Whitney test.

Principal component analysis (PCA; **Fig. 4B**) and hierarchical clustering (**Fig. 4C**) demonstrated distinct patterns of mRNA methylation in sALS vs. control sections. These studies also suggested widespread mRNA hypermethylation in sALS patient samples, with 37% of assayed m6A sites in sALS displaying increased methylation (**Fig. 4D**) whereas <1% of transcripts were hypomethylated compared to controls in sALS spinal cord. We detected a similar pattern of hypermethylation for lncRNA (**Fig. 4E, F**), indicating a broad phenomenon not limited to protein-coding mRNAs.

To determine which of the hypermethylated transcripts in ALS are m6A-modified TDP43 substrates, we examined the overlap between the DART-seq results from HEK293T cells (**Fig. 2**), and hypermethylated transcripts from sALS spinal cord (**Fig. 4G**). Within the 2184 hypermethylated transcripts that are detectable in both HEK293T cells and spinal cord, 2034 or 93% were also identified as m6A-modified RNAs by DART-seq, indicating excellent agreement between the two approaches (p∼0, hypergeometric test). These transcripts were enriched for protein kinases and RNA binding proteins, several of which are also associated with ALS (*CSNK1E, TIA1, hnRNPAB, ANXA11*), other neurodegenerative diseases (*PARK7, WDR45*), basic processes such as stress granule formation (*UBAP2L*), and nucleocytoplasmic transport (*NXF1, KPNB1*). By cross-referencing these transcripts with m6A-modified RNAs pulled-down by HaloTag-TDP43 (**Fig. 2**), we identified 302 hypermethylated TDP43-target RNAs (**Fig. 4G**) — these transcripts were likewise enriched in kinases and RNA binding proteins, and included several factors linked with ALS pathogenesis (*TP53, UCHL1*) or RNA methylation itself (*METTL3, HNRNPC*). Remarkably, the 302 hypermethylated TDP43 targets, as well as the larger set of 2034 hypermethylated transcripts in sALS spinal cord, displayed strong enrichment for RNAs whose expression is regulated by TDP43 (**Fig. 4H**). Thus, a broad range of transcripts are hypermethylated in sALS spinal cord, many of which are functionally regulated by TDP43 and overlap with m6A-modified TDP43 target RNAs highlighted by DART-seq.

As confirmation for the array, and to examine the distribution of RNA hypermethylation in sALS, we immunostained control and sALS patient spinal cord using antibodies against m6A (**Fig. 4I**). Moderate cytoplasmic staining for m6A RNA was detected in large neurons located within the anterior horn of control spinal cord. In comparison, we observed pronounced and often punctate m6A staining in anterior horn neurons from sALS sections. Indeed, m6A immunoreactivity was approximately 1.5-fold greater in sALS spinal neurons compared to controls (**Fig. 4J**). These data provide independent verification of the epitranscriptomic array (**Fig. 4C-F**) demonstrating RNA hypermethylation in sALS spinal cord, and further suggest that m6A-modified RNA accumulates predominantly within spinal motor neurons in sALS.

### YTHDF2 knockout mitigates TDP43-related neurotoxicity

Considering TDP43’s ability to recognize m6A-modified RNA, and the substantial RNA hypermethylation noted in sALS spinal cord, we questioned whether RNA methylation or the factors acting on m6A-modified RNA are involved in TDP43-mediated neurotoxicity. We therefore designed a platform that would enable us to readily screen m6A-related factors using CRISPR/Cas9 in a neuron model of ALS/FTD due to TDP43 accumulation^7,39,46,47^. As proof of principle, single-guide (sg)RNAs directed against the neuronal transcription factor NeuN or lacZ were cloned into a vector that also encodes enhanced green fluorescent protein (EGFP) and Cas9 nuclease. Rodent primary mixed cortical neurons were transfected with this vector, and after 5d, cells were fixed and immunostained for NeuN. Confirming the utility of this platform for single-cell gene knockouts, we observed a marked reduction in NeuN immunoreactivity only in transfected cells marked by EGFP fluorescence, and only in neurons that received sgRNAs against NeuN (**Fig. 5A, B**).

**Figure 5:**
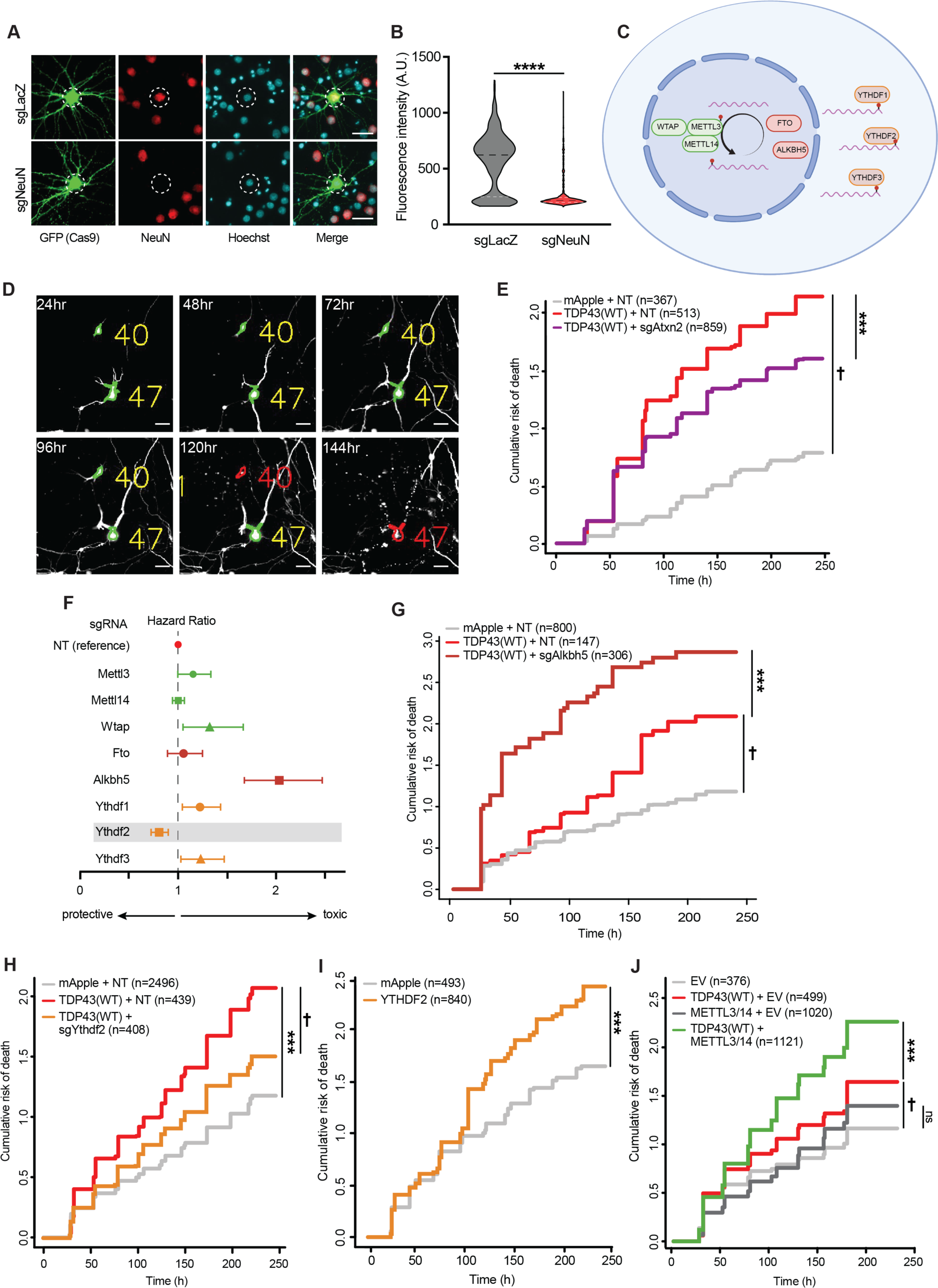
m6A factors modulate TDP43-dependent neurotoxicity. (**A**) Representative images of rodent primary neurons transfected with plasmids expressing Cas9-2A-EGFP and sgRNA targeting the neuronal protein NeuN or negative control (LacZ). 5d after transfection, neurons were fixed and immunostained for NeuN (red). White dashed circles indicate nucleus stained with Hoechst (blue). (**B**) NeuN antibody reactivity measured in EGFP-positive neurons expressing sgLacZ (n= 565) or sgNeuN (n= 654), ****p < 0.0001 by Mann-Whitney. (**C**) Schematic depicting m6A writers (green), erasers (red), and readers (orange) targeted by CRISPR/Cas9. (**D**) Primary neurons expressing EGFP and TDP43-mApple were assessed at regular 24h intervals by fluorescence microscopy, and their survival assessed by automated image analysis. Individual neurons are assigned unique identifiers (yellow number) and tracked until their time of death (red), indicated by cellular dissolution, blebbing, or neurite retraction. Scale bar= 20µm. (**E**) Cumulative hazard plot depicting risk of death for neurons expressing TDP43(WT) + non-targeting (NT) (red line), mApple + NT (grey line), or TDP43(WT) + Atxn2 sgRNA (purple line). †p<2.0 ×10^−16^, Hazard ratio (HR)= 3.45; ***p= 5.81 ×10^−4^, HR= 0.80). (**F**) Forest plot showing HR for TDP43-overexpressing neurons upon knockdown of m6A writers (green), erasers (dark red), and readers (orange), in comparison to nontargeting (NT) control. Dashed line indicates HR= 1, representing the survival of the reference condition, neurons expressing TDP43-mApple and NT sgRNA. Values >1 indicate increased toxicity, whereas values <1 denote relative protection. Error bars represent 95% CI. (**G**) Alkbh5 knockout significantly increases TDP43 associated toxicity. †p=3.11 ×10^−5^, HR= 1.59; ***p= 2.65×10^−11^, HR= 2.03. (**H**) Ythdf2 knockout significantly extends survival in TDP43-expressing neurons. ***p <2.0 ×10^−16^, HR= 1.69; †p= 6.2 ×10^−6^, HR= 0.71. (**I**) YTHDF2 overexpression is toxic to neurons. ***p= 3.07×10^−5^, HR= 1.30. (**J**) METTL3/14 overexpression enhances TDP43-dependent toxicity in neurons. †p = 5.53 ×10^−4^, HR= 1.32; ***p =4.16 ×10^−6^, HR= 1.31. p values in **E, G-J** determined via Cox proportional hazards analysis, with a minimum 3 of biological replicates.

We then designed a small-scale screen targeting the major m6A writers (METTL3, METTL14, WTAP), erasers (FTO, ALKBH5), and readers (YTHDF1, YTHDF2, YTHDF3) (**Fig. 5C**) in a neuronal model of TDP43-related disease^7,39,46,47^. sgRNAs targeting each factor were cloned into the EGFP-Cas9 vector described above and transfected into rodent primary mixed cortical neurons along with the red fluorescent protein mApple or TDP43 fused with mApple (TDP43-mApple; **Fig. 5D**). Overexpression of TDP43 recapitulates key aspects of ALS/FTD pathophysiology in neurons, including progressive neurodegeneration accompanied by TDP43 mislocalization and aggregation^48^. To track TDP43-dependent neuron loss over time, we employed automated longitudinal microscopy^49,50^. Here, hundreds of transfected neurons are simultaneously imaged at regular 24h intervals. Image segmentation algorithms detect neurons based on their unique morphology and determine their time of death through characteristic changes including soma rounding, neurite retraction and fragmentation, and cellular blebbing (**Fig. 5D**).

Transient transfection in this manner results in a wide range of expression levels, ranging from 1-7-fold endogenous TDP43 (**Sup. Fig. 5A**). As in previous studies, we observed dose-dependent TDP43-mediated toxicity in primary neurons, with the greatest risk of death observed in neurons with the highest TDP43 expression (**Sup. Fig. 5B-C**). Therefore, to maximize our ability to identify genetic modulators of TDP43 toxicity we focused on neurons within the lowest TDP43 expression (<2-fold endogenous levels), thereby avoiding supraphysiological TDP43 levels that may mask modifier effects.

In cells expressing non-targeting (NT) sgRNA, TDP43-mApple overexpression resulted in a >3-fold increase in the risk of death compared to neurons transfected with mApple alone (**Fig. 5E;** hazard ratio (HR)= 3.4; p= 2×10^−16^, Cox proportional hazards analysis). As a positive control, we cotransfected neurons with sgRNAs targeting Atxn2, one of the strongest genetic modifiers of TDP43-dependent toxicity^51–53^. Co-expression of Atxn2 sgRNA significantly suppressed TDP43-mediated toxicity (HR= 0.80; ***p= 5.81×10^−4^, Cox proportional hazards analysis), validating the use of this system for identifying disease modifiers. We then assessed whether knockout of the m6A writers, erasers or readers listed above are capable of modulating neuron loss due to TDP43 overexpression (**Fig. 5F**). In control neurons expressing mApple alone, we observed baseline increases in the risk of death upon knockdown of several factors (**Sup. Fig. 7**), suggesting that many m6A-related components are essential. Among these factors, knockout of Wtap (HR= 1.3; *p= 0.02), Alkbh5 (HR= 2.03; ***p= 2.65×10^−11^), Ythdf1 (HR= 1.23; *p= 0.01), and Ythdf3 (HR= 1.23; *p= 0.03) significantly enhanced TDP43-related toxicity (**Fig. 5G** and **Sup. Figs. 6**,**7**). Only Ythdf2 knockout (HR= 0.71; ***p= 6.2×10^−6^) reduced TDP43-mediated toxicity (**Fig. 5H**), suggesting that TDP43 may act in concert with YTHDF2 to elicit neurodegeneration.

In a complementary set of experiments, we overexpressed a subset of m6A-related factors in rodent primary neurons and followed neuronal survival by automated longitudinal microscopy. Analogous to TDP43, overexpression of YTHDF2 resulted in a significant increase in the risk of death compared to the negative control (**Fig. 5I**). Conversely, overexpression of METTL3 and METTL14 had little effect on their own in primary neurons (**Fig. 5J**). To determine if METTL3/METTL14 might act synergistically with TDP43, we expressed TDP43-mApple at low concentrations, resulting in more subtle toxicity than in previous experiments. In doing so, we found that METTL3/METTL14 overexpression significantly enhanced TDP43-related toxicity (**Fig. 5J**). Considering the ability of TDP43 to bind m6A-modified RNA, and the increase in RNA methylation stimulated by METTL3/METTL14 overexpression^54^, these data imply that RNA hypermethylation facilitates TDP43-related neurodegeneration in ALS/FTD models.

### YTHDF2 in sALS spinal cord and human iPSC-derived neurons

We detected relative RNA hypermethylation in sALS spinal cord samples by m6A array (**Fig. 4**) and found that genetic ablation of the m6A reader YTHDF2 reduced TDP43-dependent toxicity in a neuronal model of disease (**Fig. 5H**). Furthermore, YTHDF2 overexpression was sufficient to elicit toxicity in rodent primary neurons (**Fig. 5I**). To determine if YTHDF2, like TDP43, is mislocalized and/or accumulates in ALS, we performed immunohistochemistry for YTHDF2 in human post-mortem samples from sALS and control spinal cord (**Fig. 6A, B**) and frontal cortex (**Sup. Fig. 8**). In controls, YTHDF2 displayed the expected uniform, cytoplasmic distribution within neurons and other cell types. Not only was YTHDF2 staining significantly more intense within sALS spinal neurons, but we also noted punctate accumulations of YTHDF2 within many of these cells (**Fig. 6A, B**). Strong YTHDF2 staining was also noted within layer IV-V neurons from ALS frontal cortex (**Sup. Fig. 8**), but puncta were not as evident in these cells as in spinal motor neurons. These results show that YTHDF2 accumulates in association with RNA hypermethylation in sALS spinal neurons, potentially contributing to TDP43-mediated toxicity and disease pathogenesis.

**Figure 6:**
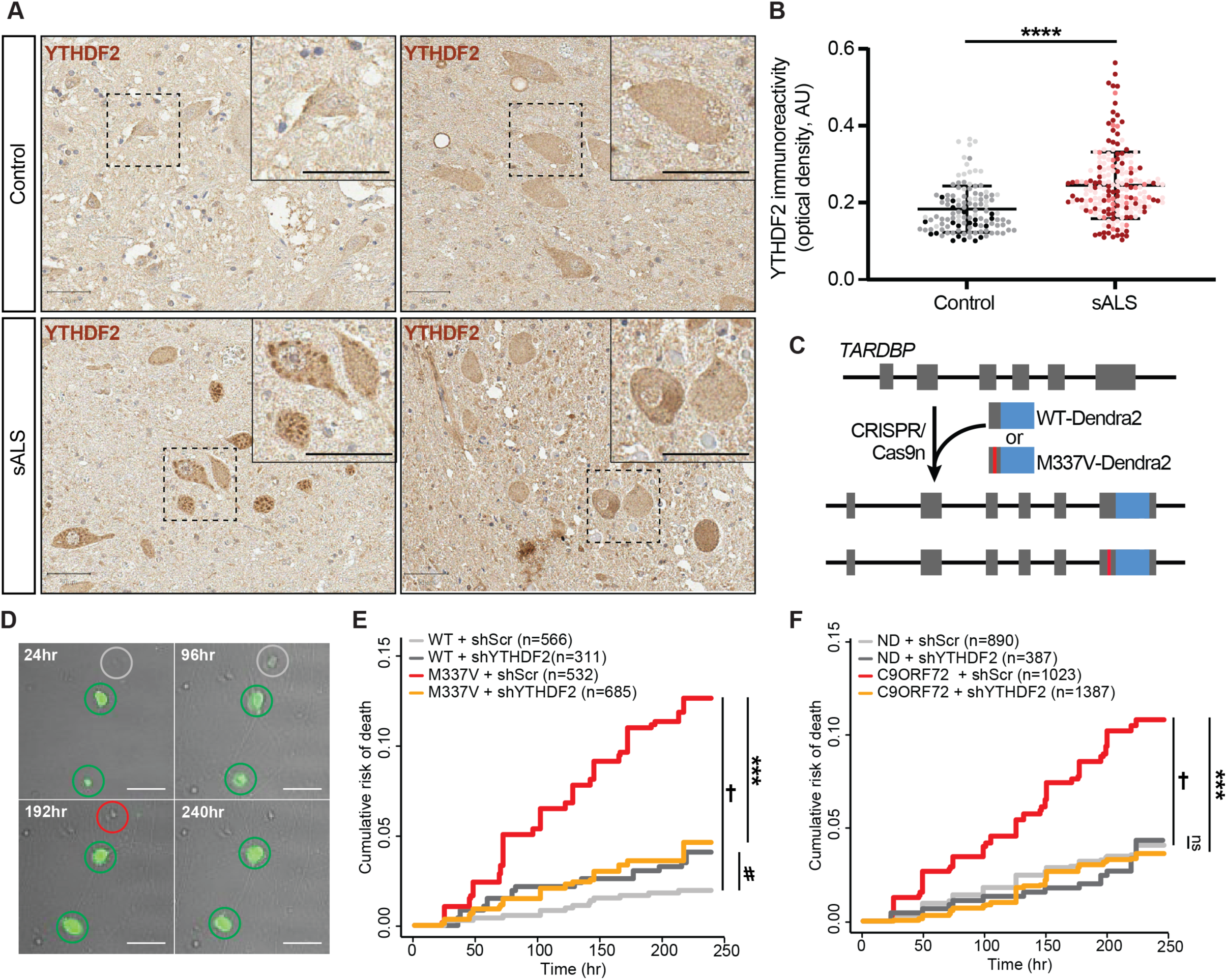
YTHDF2 reduction extends neuronal survival in human neuron disease models. (**A**) Immunostaining of YTHDF2 in control and sALS patient spinal cord samples. Scale bar= 50 µm. (**B**) Quantification of YTHDF2 immunoreactivity in spinal cord neurons from control (n= 117 neurons) and sALS (n= 193 neurons) samples. Plot shows mean +/- SD, color coded by sample. ****p< 0.0001 via Mann-Whitney test. (**C**) Strategy used to create isogenic iPSCs expressing native TDP43(WT)-Dendra2 or TDP43(M337V)-Dendra2. (**D**) Representative images of untransduced (grey) and transduced (green) iNeurons expressing shRNA against YTHDF2 (shYTHDF2) and a GFP reporter. Time of death (red circles) for each cell is used to determine cumulative risk of death, plotted in (**E**) and (**F**). Scale bar= 20µm. shRNA-mediated knockdown of YTHDF2 significantly extended the survival of TDP43(M337V)-Dendra2 iNeurons (**E**; †p= 8.42×10^−12^, HR= 6.25; ***p= 4.82×10^−9^, HR=0.32; #p= 0.08, HR= 1.84) as well as mutant *C9ORF72* iNeurons (**F**, †p= 1.42×10^−11^, HR= 2.85; ***p= 1.42×10^−16^, HR= 0.32). ns= not significant. Values in (**E**,**F**) calculated by Cox proportional hazards analysis, with a minimum 3 biological replicates.

The overabundance of YTHDF2 in sALS spinal cord, together with data from **Fig. 5** suggesting neuroprotection upon Ythdf2 knockout in primary neurons, imply that YTHDF2 could be a therapeutic target for sALS. To pursue this further, we examined the effect of *YTHDF2* knockdown in human neuron disease models. The ALS-associated TDP43(M337V) mutation was introduced into the native *TARDBP* locus of control human induced pluripotent stem cells (iPSCs) by CRISPR/Cas9^55^. Simultaneously, endogenous TDP43 was labeled at the C-terminus with Dendra2, facilitating identification and selection of clonal cell populations harboring the M337V mutation. We also generated isogenic WT iPSCs in which native TDP43 was fused to Dendra2 without introduction of a pathogenic mutation (**Fig. 6C**). iPSCs were then differentiated into forebrain-like neurons (iNeurons) through the induced expression of master transcription factors Ngn1-2, as described previously^40,55,56^. Neurons were tracked by longitudinal microscopy over a 10d period, and their survival assessed by semi-automated tracking software developed specifically for these purposes (**Fig. 6D**). We detected a significant increase in the risk of death for TDP43(M337V) iNeurons in comparison to controls (**Fig. 6E**, †p= 8.42×10^−12^, HR= 6.25; Cox proportional hazards analysis) after sustained growth in the same culture after differentiation (see methods). As in our primary neuron disease model, *YTHDF2* knockdown via lentiviral shRNA delivery substantially extended the survival of TDP43(M337V) iNeurons without adversely affecting WT iNeurons (**Fig. 6E**, ***p= 4.82×10^−9^, HR= 0.32; #p= 0.08, HR= 1.84; Cox proportional hazards analysis).

We also evaluated the effect of *YTHDF2* knockdown in 3 separate lines of human iNeurons carrying the *C9ORF72* hexanucleotide expansion, the most prevalent cause of familial ALS and FTD in Northern Europe and North America^57–59^. *C9ORF72* mutant iNeurons displayed a ∼3-fold elevation in the risk of death upon neurotrophic factor withdrawal compared to control neurons (**Fig. 6F**, †p= 1.42×10^−11^, HR= 2.86; Cox proportional hazards analysis). As in TDP43(M337V) iNeurons, shRNA-mediated knockdown of *YTHDF2* significantly prolonged the survival of *C9ORF72* mutant iNeurons but had no effect on non-disease (ND) iNeurons (***p= 1.42×10^−16^, HR= 0.32; Cox proportional hazards analysis). Together with our data from rodent primary neurons (**Fig. 5**), these findings emphasize the neuroprotective potential of *YTHDF2* knockdown in ALS/FTD disease models featuring TDP43 pathology.

## Discussion

Here, we show not only that TDP43 recognizes m6A-modified RNA, but also that the majority of TDP43 substrates exhibit m6A marks. Sequence analysis demonstrated a spatial correlation between 3’UTR m6A modifications and TDP43 recognition motifs, and further showed that methylation strongly influences the binding of TDP43 to its RNA targets as well as TDP43 autoregulation. In ALS, where cytoplasmic mislocalization and aggregation of TDP43 are signature pathologic changes, we also observed widespread RNA hypermethylation compared to disease controls. Consistent with a primary role for RNA hypermethylation in ALS pathogenesis, knockout or knockdown of the m6A writer YTHDF2 mitigated toxicity in primary rodent and human iPSC-derived neuron models of ALS and FTD, while overexpression of the m6A writers METTL3 and METTL14 exacerbated TDP43-mediated neuron loss. Together, these findings underscore the importance of m6A modifications for RNA binding by TDP43 and emphasize the potential contribution of TDP43’s actions on m6A-modified RNA to the development of ALS and FTD.

Our study builds on several lines of evidence hinting at a connection between TDP43 and m6A-modified RNA. m6A writers (METTL3, METTL14, WTAP) and erasers (FTO, ALKBH5) primarily localize to nuclear speckles^60,61^, membraneless organelles that facilitate peritranscriptional RNA processing and splicing. TDP43 is also concentrated within these structures^62^, and interacts with several m6A writer and reader proteins, including METTL3, YTHDF1, YTHDF2 and hnRNPC^15,63,64^. Furthermore, TDP43 was among a series of proteins that exhibited preferential binding to m6A-modified bait RNAs^65^. Supporting this, we found that both overexpressed and endogenously labeled TDP43 in HEK293T cells were capable of pulling down m6A modified RNA. Combining affinity purification of native HaloTag-TDP43 with antibody-free detection of m6A modifications through DART-seq, we were able to clarify the location and number of m6A sites within TDP43 substrate RNAs. Importantly, DART-seq is not limited by the specificity or sensitivity of antibodies for m6A modifications, instead relying on APOBEC1-YTH induced C-T transitions to indicate possible m6A sites. In contrast to the previously noted enrichment of m6A marks within the proximal 3’UTR of transcripts^11,12,25,27^, TDP43 targets displayed a broad distribution of m6A modifications across the CDS and 3’UTR. It is unclear whether this pattern is unique to TDP43 substrate RNAs, or whether a similar distribution of m6A marks would be observed in targets of related RNA binding proteins. We observed few m6A sites within the introns of TDP43 target RNAs, consistent with the relative lack of intronic m6A modifications noted in previous studies^66–68^. Given the relative concentration of TDP43 binding sites within introns and 3’UTRs^36,37^, and our finding that m6A modifications cluster within the CDS and 3’UTR of TDP43 target RNAs, it stands to reason that the overlap between TDP43 binding sites and m6A modifications is greatest within the 3’UTR of these transcripts. We anticipate that the expression and/or stability of this subset of TDP43 target RNAs is strongly influenced by conditions that enhance or reduce m6A modifications.

To assess the consequences of such m6A modifications, we focused on the *TARDBP* transcript itself. Previous studies as well as our own DART-seq experiments identified key m6A modifications located within the TBR, a region of the *TARBDP* transcript that is essential for recognition and autoregulation by TDP43 itself^38,42^. In keeping with the association between m6A modifications and UG-rich domains recognized by TDP43 (**Fig. 1**), these m6A marks are located immediately adjacent to TDP43 binding motifs within the TBR. Mutagenesis of the methylated A within a reporter containing the TBR led to a reduction in TDP43 binding and ineffective autoregulation, suggesting that m6A modifications are crucial for TDP43 recognition and events downstream of binding. Even so, *in vitro* electromobility shift assays involving recombinant TDP43 and a short (14 nt) m6A-modified probe failed to show methylation-dependent increases in TDP43 binding, implying that m6A modifications indirectly enhance RNA recognition by TDP43. Similar relationships have been observed for hnRNP-C^20^ and IGF2BP3^69^, suggesting that changes in RNA secondary or tertiary structure upon methylation can promote recognition of otherwise buried motifs by TDP43 and other RNA binding proteins.

In prior work, we observed extensive RNA destabilization in human iPSCs overexpressing TDP43^5^. Transcripts encoding components of the ribosome and oxidative phosphorylation pathways were most heavily represented among TDP43-destabilized RNAs, a pattern that was mirrored in iPSCs carrying ALS/FTD-associated *C9ORF72* mutations. Notably, these same families of transcripts are upregulated upon targeted deletion of the m6A writer METTL14^10^, indicating that they are regulated by m6A modification. Given these observations, together with the degree of RNA hypermethylation we observed in sALS spinal cord and the ability of TDP43 to recognize m6A-modified RNA, we suspect that TDP43 mislocalization and accumulation may lead to RNA destabilization in ALS through an m6A-dependent mechanism. Although this hypothesis requires further investigation, as an example we explored the potential for m6A modifications to influence TDP43 autoregulation, a phenomenon that involves the nuclear retention and destabilization of *TARDBP* transcripts upon their recognition by TDP43^5,36,43–45^. m6A marks located within the TBR influence TDP43 binding and are crucial for proper autoregulation of the protein. In light of recent data highlighting the possible contribution of TDP43 autoregulation to ALS/FTD pathogenesis^45,70^, these observations draw attention to the potential importance of m6A modifications for physiological TDP43 function as well as its dysfunction in disease.

Among different tissue types, the central nervous system displays some of the highest baseline levels of m6A RNA, and these modifications are critical for proper neuronal development and maturation^12,71–73^. Total m6A levels within the nervous system rise with age, and previous evidence suggests that neurodegenerative diseases independent of ALS and FTD are likewise characterized by RNA hypermethylation^74,75^. We observed significant hypermethylation in end-stage tissue from ALS patients compared to age-matched controls; although we do not believe age is responsible for the observed RNA hypermethylation, other factors may be playing a part. Since astrocytosis and microgliosis are common features of ALS as well as other neurodegenerative diseases, it is possible that much of the observed RNA hypermethylation arises not because of a primary disease mechanism, but instead because of secondary neuroinflammation^76^.

Importantly, YTHDF2 was also the only m6A-related factor that emerged from our limited CRISPR-based screen for modulators of TDP43-mediated toxicity. This platform, which combines longitudinal fluorescence microscopy, automated survival analysis, and single-cell gene knockout, allowed us to rapidly interrogate most m6A readers, writers, and erasers for their effects on neuronal survival. In doing so, we found that knockout or knockdown of the m6A reader *Ythdf2* mitigated TDP43-related toxicity in rodent primary neurons, while YTHDF2 overexpression itself was lethal, and transfection with writers (METTL3 and METTL14) exacerbated TDP43-dependent neuron loss. Furthermore, *YTHDF2* knockdown in human iNeurons carrying ALS/FTD-associated mutations in *TARDBP* and *C9ORF72* prolonged cellular survival. These data, together with the apparent accumulation of YTHDF2 in ALS spinal cord, suggest that YTHDF2 may be a novel therapeutic target in ALS and FTD.

## Methods

### HEK293T cell culture

Human embryonic kidney (HEK) 293T cells were cultured in DMEM (GIBCO), 10% FBS, 100 units/mL Penicillin/Streptomycin at 37°C in 5% CO2. HEK293T cells are originally female in origin, are easily transfected, and have been transformed with SV40 T-antigen.

### CRISPR/Cas9 integration of HaloTag into *TARDBP* locus in HEK293T cells

Oligos complementary to the target region (**Table S4**) were annealed, digested, and ligated into the *BbsI* site in the pX335 vector (Addgene, #42335) according to the protocol available from Addgene. 1.25 µg of each vector was transfected into HEK293T cells using Lipofectamine 2000 (ThermoFisher, #11668019) together with 2.5 µg of a plasmid encoding the HaloTag open reading frame flanked by 400 bp of sequence homologous to regions immediately upstream and downstream of the *TARDBP* start codon, according to the manufacturer’s instructions. Following transfection, cells were split at a low density, allowing transfected cells to establish individual colonies. Cells were screened for nuclear fluorescence after incubation with JF635 dye, as described previously^40,77^. Positive cells were carefully scraped/aspirated using a P200 pipet tip and transferred to a new dish. This process was repeated until 100% of cells displayed nuclear fluorescence after JF635 application, and correct integration of the HaloTag cassette into the TARDBP locus was confirmed by PCR and Sanger sequencing. HaloTag-TDP43 HEK293T cells were cultured on plates coated with 0.1% gelatin (Sigma, #G2500) in DMEM (GIBCO), 10% FBS, 100 units/mL Penicillin/Streptomycin at 37°C in 5% CO2.

### iNeuron differentiation

Day 0: Induced pluripotent stem cells were washed in PBS and incubated in prewarmed accutase (Sigma, #A6964) at 37°C for 8 min. Four volumes of E8 media (ThermoFisher, #A1517001) were added to the plate, and the cells were collected and pelleted at 200xg for 5 min. The media was aspirated, and the pellet was resuspended in 1mL of fresh E8 media. Cells were counted using a hemocytometer, diluted, plated at a density of 20,000 cells/mL in E8 media with ROCK inhibitor and incubated at 37°C overnight. Day 1: Media was changed to N2 media (1x N2 Supplement (Gibco, #17502-048), 1x NEAA Supplement (Gibco, #11140-050), 10 ng/mL BDNF (Peprotech, #450-02), 10 ng/mL NT3 (Peprotech, #450-03), 0.2 µg/mL laminin (Sigma, #L2020), 2 mg/mL doxycycline (Sigma, #D3447) in E8 media). Day 2: Media was changed to transition media ((1x N2 Supplement, 1x NEAA Supplement, 10 ng/mL BDNF, 10 ng/ml NT3, 0.2 µg/mL laminin, 2 mg/mL doxycycline in half E8 media, half DMEM F12 (Gibco, #11320-033)). Day 3: Media was changed into B27 media (1x B27 Supplement (Gibco, #17504-044), 1x Glutamax Supplement (Gibco, #35050-061), 10 ng/mL BDNF, 10 ng/mL NT3, 0.2 µg/mL laminin, and 1x Culture One (Gibco, #A33202-01) in Neurobasal-A (Gibco, #12349-015)). On day 6 cells were transduced with the appropriate virus (prepared by University of Michigan Vector Core, **Table S2**) and cells were sustained in the same culture medium for the remainder of the experiment. Day 14: Imaging began for survival experiments and iNeurons were monitored over the course of 10 days.

### Plasmids

pGW1-GFP, pGW1-TDP43(WT)-GFP, pGW1-mApple^46,48^, pGW1-Halo, pGW1-TDP43(WT)-Halo^7,40^, and pCAGGS-TDPBR-mCherry^7,39,40^ were created as previously described. To generate pGW1-YTHDF2-Halo, the YTHDF2 ORF was PCR amplified from pcDNA-flag-YTHDF2 (Addgene, #52300). The resulting amplicon was digested with AgeI and XbaI and cloned into cut sites upstream of HaloTag insert in pGW1-Halo.

To create pgW1-YTHDF2-2A-GFP, the YTHDF2 ORF was PCR amplified from pcDNA-flag-YTHDF2 (Addgene, #52300). The resulting amplicon was digested with KpnI and SalI and cloned into cut sites upstream of the 2A in the pGW1-2A-GFP vector.

pCAGGS-TDPBR (mTBR)-mCherry, was created via site directed mutagenesis from pCAGGS-TDPBR-mCherry using the Pfu Ultra high-fidelity polymerase (Agilent Technologies, #600380) according to manufacturer’s protocols to change the A into a G in clip34nt sequence in the TDPBR.

### HaloTag immunoprecipitation

HEK293T cells were transfected with pGW1-Halo, pGW1-TDP43-Halo, or pGW1-YTHDF2-Halo using Lipofectamine 2000 (ThermoFisher, #11668019) following the manufacturer’s protocol. Approximately forty-eight hours post-transfection, cell pellets were harvested and lysed using a syringe in 100 µL lysis buffer (50 mM Tris-HCl, 150 mm NaCl, 1% Triton-X100, 0.1% NaDeoxycholate) and incubated on ice for 30 min. Following lysis, tubes were centrifuged at 17000xg for 5 min and pellet discarded. 200 g for lysate was added to 25 µL of HaloTrap beads (Chromotek, #ota-10), prewashed with 500 µL of 10 mM Tris-HCl (pH 7.5), 150 mM NaCl, 0.5 mM EDTA centrifuged at 2500xg for 5 minutes. Lysate and bead mix was incubated rotating overnight at 4°C. Following incubation, the mixture was centrifuged at 2500xg for 5 min and supernatant discarded. HaloTrap beads were washed 3 times for 5 min at 2500xg in 500 µL of wash buffer (50 mM Tris-HCl, 150 mM NaCl, 1% Triton-X100, 0.1% NaDeoxycholate, 0.05% IGEPAL). To elute RNA, Trizol (ThermoFisher, #15596026) was added directly to the beads and RNA was isolated using phenol-chloroform extraction.

### m6A dot blot

For m6A dot blot, isolated RNA was boiled at 95°C for 3 minutes to denature RNA then immediately chilled on ice. RNA was added in 1 µL drops to a BrightStar nylon membrane (ThermoFisher, #AM10102) and crosslinked at 1200 µJ [x100] for 2 min twice. After crosslinking, membranes were stained with methylene blue (0.04% methylene blue in 0.5M sodium acetate pH 5.2) until circles were visible (approximately 5 min). Membranes were washed 3x with water quickly to remove background stain and imaged. Afterwards, membranes were washed several times with water to remove methylene blue staining, then blocked in 5% milk in 1x PBS + 0.02% Tween-20 at room temperature for 1 hour. Primary antibody (rabbit anti-m6A antibody 1:500, Cell Signaling #56593) was added to blocking solution, and membranes rocked overnight at 4°C. Blots were then washed in wash buffer (1x PBS + 0.02% Tween-20) 3x for 5 min and incubated at room temperature with blocking buffer containing goat anti-rabbit HRP secondary antibody (Jackson Immunoresearch labs, #111-035-003, 1:5000). Blots were rinsed in wash buffer 3x for 5 min, then incubated in Pierce ECL Western blotting substrate (ThermoFisher, #32106) for 1 min before exposure to CL-XPosure™ Film (Fisher, #PI34090). For endogenous Halo-TDP43 pulldowns, the procedure was followed as before starting from cell pellets of Halo-TDP43 HEK293T or unmodified HEK293T cells.

### DART-seq

Halo-TDP43 HEK293T cells were transfected with pCMV-APOBEC1-YTH, pCMV-APOBEC1-YTHmut, or pGW1-Halo using Lipofectamine 2000 (ThermoFisher, #11668027) following the manufacturer’s protocol. Approximately 48 hours post-transfection, cells were washed with cold PBS (Invitrogen) then crosslinked with UV light (254 nm, 150 mJ/cm^2^). Following crosslinking, cells were harvested and lysed using a syringe in 100 µL lysis buffer supplemented RNAse inhibitor (Invitrogen, #N8080119) (50 mM Tris-HCl, 150 mM NaCl, 1% Triton-X100, 0.1% NaDeoxycholate) and incubated on ice for 30 min. 200 µg for lysate was added to 25 µL of HaloTrap beads, prewashed with 500 µL of equilibration buffer supplemented with RNAse inhibitor (10 mM Tris-HCl (pH 7.5), 150 mM NaCl, 0.5 mM EDTA) centrifuged at 2500xg for 5 minutes. Lysate and bead mix was incubated rotating overnight at 4°C with Baseline Zero DNAse (Fisher Scientific, #NC1424104). Following incubation, the mixture was centrifuged at 2500xg for 5 min and supernatant discarded. HaloTrap beads were washed 3 times for 5 min at 2500xg in 500 µL of wash buffer supplemented with RNAse inhibitor (50 mM Tris-HCl, 150 mM NaCl, 1% Triton-X100, 0.1% NaDeoxycholate, 0.05% IGEPAL). To elute RNA, Trizol was added directly to the beads and RNA was isolated using phenol-chloroform extraction. Once extracted, RNA was submitted to Advanced Genomics Core at University of Michigan for RNA sequencing.

### Next-generation sequencing

cDNA libraries were prepared from Trizol extracted, DNA-digested samples using the Illumina Stranded Total RNA Prep with Ribo-Zero Plus kit (Illumina, #20040525). Paired end sequencing was carried out on an Illumina NovaSeq (S4) 300 cycle sequencer at the University of Michigan Advanced Genomics Core. Samples were sequenced at an average depth of 15.6 million unique reads per sample.

### DART-seq data processing and analysis

Raw reads were quality and adapter trimmed with Cutadapt (v2.3)^78^ with default parameters, and aligned to the GRCh38 human genome assembly with bwa-mem (v0.7.17)^79^ with default parameters. Single nucleotide transition analysis was performed as previously described^27^. Briefly, aligned reads were deduplicated, sorted by genome coordinate, parsed to BED format, and collapsed by PCR replicate using the CLIP Tool Kit (CTK) suite^80^. Mutations were extracted and replicates were merged, preserving replicate information. The CTK suite script “CIMS.pl” was then used to generate coverage, mutation rate, and a false discovery statistic. Resulting BED files were filtered in R for entries with a minimum mutation rate of 2, a minimum read count of 10 per replicate, and a mutation/read threshold of 0.1-0.6. Finally, sense C to T (and antisense G to A) transitions were filtered for presence of the surrounding DRACH motif using custom scripts and the GenomicRanges, BSgenome, Biostrings, and gUtils R packages.

The distribution of mutational transitions was calculated using custom scripts adapted from the MetaPlotR Perl/R suite^81^ and visualized with ggplot2. Motif logos were generated before and after DRACH filtration using ggseqlogo. Linear U-G dinucleotide density was calculated using custom R scripts. Briefly, sequences were collapsed to binary representations of UG/GU (1) or non-UG/GU (0) dyads. For UG15 density, a continuous sliding average was calculated in 15-nucleotide windows along the target sequence such that a tract of uniform UG alternation corresponds to a UG15 density of 1. Sliding averages were rescaled to the basepair length of the target sequence and used to generate site-of-interest-centered density plots using ggplot2. Gene ontology analyses were accomplished via STRING or Enrichr^29,82^. Euler diagrams were created using eulerr in R^83^.

### m6A array

Human spinal cord samples were homogenized in Trizol and RNA was extracted using phenol-chloroform extraction for the m6A mRNA&lncRNA Epitranscriptomic microarray (8×60K, Arraystar, Rockville, MD, USA). 1-3 μg total RNA and m6A spike-in control mixture were added to 300 μL 1xIP buffer (50 mM Tris-HCl, pH 7.4, 150 mM NaCl, 0.1% NP40, 40U/μL RNase Inhibitor) containing 2 μg anti-m6A rabbit polyclonal antibody (Synaptic Systems, #202003). The reaction was incubated with head-over-tail rotation at 4°C for 2 hours. 20 uL DynabeadsTM M-280 Sheep Anti-Rabbit IgG (Invitrogen, #11203D) suspension per sample was blocked with freshly prepared 0.5% BSA at 4°C for 2 hours, washed three times with 300 μL 1xIP buffer, and resuspended in the total RNA-antibody mixture prepared above. The RNA binding to the m6A-antibody beads was carried out with head-over-tail rotation at 4°C for 2 hours. The beads were then washed three times with 500 μL 1xIP buffer and twice with 500 μL Wash buffer (50 mM Tris-HCl, pH7.4, 50 mM NaCl, 0.1% NP40, 40 U/μL RNase Inhibitor (Enzymatics, #Y9240L)). The enriched RNA was eluted with 200 μL Elution buffer (10 mM Tris-HCl, pH7.4, 1 mM EDTA, 0.05% SDS, 40U Proteinase K) at 50°C for 1 hour. The RNA was extracted by acid phenol-chloroform and ethanol precipitated. The “IP” RNA and “Sup” RNAs were added with equal amount of calibration spike-in control RNA, separately amplified and labeled with Cy3 (for “Sup”) and Cy5 (for “IP”) using Arraystar Super RNA Labeling Kit (Arraystar, #AL-SE-005). The synthesized cRNAs were purified by RNeasy Mini Kit (QIAGEN, #74105). The concentration and specific activity (pmol dye/μg cRNA) were measured with NanoDrop ND-1000. 2.5 μg of Cy3 and Cy5 labeled cRNAs were mixed. The cRNA mixture was fragmented by adding 5 μL 10x Blocking Agent and 1 μL of 25x Fragmentation Buffer, heated at 60°C for 30 min, and combined with 25 μL 2x Hybridization buffer. 50 μL hybridization solution was dispensed into the gasket slide and assembled to the m6A-mRNA & lncRNA Epitranscriptomic Microarray slide. The slides were incubated at 65°C for 17 hours in an Agilent Hybridization Oven. The hybridized arrays were washed, fixed and scanned using an Agilent Scanner G2505C. Comparisons with DART-seq results (Fig. 4G) included a limited set of transcripts (2184 out of 5646) that were expressed in both human spinal cord and HEK293T cells [normalized transcripts per million (TPM) > 2], based on datasets made available through the Human Protein Atlas.

### Longitudinal microscopy and automated survival analysis

Cortices from embryonic day (E)19-20 Long-Evans rat embryos were dissected and disassociated, and primary neurons were plated at a density of 6×10^5^ cells/ml in 96-well plates, as described previously^7,39,46,84^. For CRISPR candidate screen, on *in vitro* day (DIV) 4, neurons were transfected with 25 ng of plasmids containing sgRNAs for respective genes, 50 ng of pGW1-mApple or pGW1-TDP43-mApple, and 25 ng of pGW1-EGFP to mark cell bodies using Lipofectamine 2000 (Invitrogen #52887. Following transfection, cells were placed in Neurobasal Complete Media (Neurobasal (Gibco, #21103-049), 1x B27, 1x Glutamax, 100 units/mL Pen Strep (Gibco, #15140-122)) and incubated at 37°C in 5% CO2.

### m6A CRISPR candidate screen

Oligos complementary to m6A pathway component genes were created using ChopChop^85–87^ (https://chopchop.cbu.uib.no/) then annealed, digested, and ligated into the *BbsI* site of pSpCas9(BB)-2A-GFP plasmid (Addgene, #48138) according to manufacturer’s protocol. 25 ng of plasmids containing sgRNAs for respective genes were transfected into rat primary neurons on DIV 4 using Lipofectamine 2000 (ThermoFisher, #11668027) along with 50 ng of TDP43(WT)-EGFP or mApple then survival was measured over the course of 10 days. Outcomes were calculated based on Cox proportional hazard values and related to expression of TDP43(WT)-EGFP to determine if knockout of target was beneficial or toxic.

Neurons were imaged as described previously^7,39,46,84^ using a Nikon Eclipse Ti inverted microscope with PerfectFocus3a 20X objective lens and either an Andor iXon3 897 EMCCD camera or Andor Zyla4.2 (+) sCMOS camera. A Lambda 421 multi-LED light source (Sutter) with 5 mm liquid light guide (Sutter) was used to illuminate samples, and custom scripts written in Beanshell for use in µManager controlled all stage movements, shutters, and filters. Custom ImageJ/Fiji macros and Python scripts were used to identify neurons and draw both cellular and nuclear regions of interest (ROIs) based upon size, morphology, and fluorescence intensity. Fluorescence intensity of labeled proteins was used to determine protein localization or abundance. Custom Python scripts were used to track ROIs over time, and cell death marked a set of criteria that include rounding of the soma, loss of fluorescence and degeneration of neuronal processes.

### Immunocytochemistry

HEK293T cells were plated on glass coverslips (Fisher Scientific, #1254580) and grown overnight at 37°C with 5% CO_2_. The following day, JF646 (Promega, #GA1120) was added to HEK293T media at 1:10,000 dilution and added to cells for 30 minutes. After 30 minutes, JF646 containing media was removed and washed twice with regular HEK293T media for 15 minutes each. Following wash out, coverslips were washed once with 1x PBS (Gibco, #14200-075) and fixed in 4% PFA for 10 minutes. Cells were then permeabilized with PBS + 0.1% Triton-X-100 for 20 minutes at room temperature, treated with 10 mM glycine for 20 minutes at room temperature, and then blocked in blocking buffer (PBS + 0.1% Triton-X-100, 2% fetal calf serum, and 3% BSA) for 1 hour at room temperature. Coverslips were then incubated overnight with blocking buffer + rabbit anti-TDP43 antibody (Proteintech, #10782-2-AP) at 1:500 to stain for TDP43. Following primary incubation, coverslips were washed three times with PBS for 5 minutes then stained with secondary antibody goat anti-rabbit AF488 (ThermoFisher, #A-11008) at 1:250 for 1 hour at room temperature in PBS + 0.1% Triton-X-100, 2% fetal calf serum, and 3% BSA. After secondary incubation, coverslips were washed three times for 5 minutes in PBS then stained with Hoechst 1:20,000 to mark nucleus. The coverslips were mounted on coverslips and allowed to dry overnight before imaging. Coverslips were imaged using confocal mode on ONI Nanoimager.

### Immunohistochemistry

Immunostaining was accomplished using the Dako Autostainer Link 48 (Agilent, USA). Anti-YTHDF2 antibody (Proteintech, #24744-1-AP, 1:300) or anti-m6A antibody (Synaptic Systems #202003, 1:100) were used with the Dako High pH Target Retrieval Solution (Tris/EDTA, pH 9; Agilent, USA) (20 minutes, 97°C) and the Dako Envision Flex Plus Mouse Link Kit (Agilent, USA) to detect the antibody along with the Dako DAB (Agilent, USA). Whole-slide images were generated by the University of Michigan Digital Pathology group within the Department of Pathology using a Leica Biosystems Aperio AT2 scanner equipped with a 0.75 NA Plan Apo 20x objective; 40x scanning is achieved using a 2x optical magnification changer. Resolution is 0.25 microns per pixel for 40x scans. Focus during the scan is maintained using a triangulated focus map built from individual focus points determined in a separate step before scanning is started. Proprietary software is used for image processing during acquisition. Image analysis performed using QuPath software^88^ and mean intensity calculated for stained cells.

### RT-PCR and quantitative RT-PCR

Total RNA was extracted using Trizol following the manufacturer’s protocol. To synthesize cDNA, 500 ng of total RNA was used in a 20 µL reaction volume with the Bio-Rad iScript cDNA synthesis kit (Bio-Rad, #1708890) according to the manufacturer’s protocol. The reactions were incubated at 25°C for 5 min, 46°C for 20 min, and 95°C for 1 min. For quantitative RT-PCR (qRT-PCR), reactions were carried out using Step One Plus Realtime PCR system (Applied Biosystems). Reactions were carried out using PowerUp™ SYBR™ Green Master Mix (ThermoFisher #A25742), with 1 µM primers, and 1 µL cDNA, according to manufacturer’s protocol. Relative gene expression was calculated using the ΔΔCt method. Values obtained from qRT-PCR were plotted in GraphPad Prism.

### Statistical Analysis

Statistical analysis performed using GraphPad Prism 9 or Superplots^89,90^. For primary neuron survival analysis, the open-source R survival package was used to determine hazard ratios and statistical significance between conditions through Cox proportional hazards analysis^7,40,50,91^.

### Ethics statement

All vertebrate animal work was approved by the Committee on the Use and Care of Animals (UCUCA) at the University of Michigan. All experiments were performed in accordance with UCUCA guidelines and designed to minimize animal use. Rats (*Rattus norvegicus*) were housed single in chambers equipped with environmental enrichment and cared for by veterinarians from the Unit for Laboratory Animal Medicine at the University of Michigan. All individuals were trained and approved in the care of long-term maintenance of rodents, in accordance with the NIH-supported Guide for the Care and Use of Laboratory Animals. All personnel handling the rats and administering euthanasia were properly trained in accordance with the University of Michigan Policy for Education and Training of Animal Care and Use Personnel. Euthanasia followed the recommendations of the Guidelines on Euthanasia of the American Veterinary Medical Association. Brains from individual pups in each litter were pooled to maximize cell counts prior to plating; as a result, primary cortical neurons used for all studies include an even mix of cells from both male and female pups.

## Acknowledgements

We thank the patients that donated tissue samples to make this work possible. Skin samples from the study participants were collected and de-identified in collaboration with the Michigan Institute for Clinical and Health Research (MICHR, UL1TR000433) through an institutional review board (IRB)-approved protocol (HUM00028826). We thank Dr. Stephen A. Goutman, Director of the University of Michigan ALS Clinic and Biorepository, and Crystal Pacut from the Program for Neurology Research and Discovery. We also would like to acknowledge Mr. Matthew D. Perkins for his help with postmortem tissue from the University of Michigan Brain Bank, Kathy Toy for her expertise with immunohistochemistry, John Moran for assistance with primary neuron survival experiments, and Drs. A. Malik and K. Weskamp for their suggestions.

This work was supported by National Institutes of Health (R01NS097542 and R01NS113943 to SJB; and P30AG072931 to the University of Michigan Brain Bank and Alzheimer’s Disease Research Center), the family of Angela Dobson and Lyndon Welch, the A.\ Alfred Taubman Medical Research Institute, the Danto Family, Ann Arbor Active Against ALS, and the Robert Packard Center for ALS Research. Immunohistochemistry was performed at the Rogel Cancer Center Tissue and Molecular Pathology Shared Resource Laboratory at the University of Michigan (NIH P30 CA04659229).

## Author Contributions

S.J.B. and M.M. designed the study. M.M. performed dot blots, immunoprecipitations, qRT-PCR, site directed mutagenesis, transfections, fluorescence microscopy, primary neuron survival experiments, electromobility shift assays, RNA isolations for DART-seq and the epitranscriptomic array, and immunohistochemical quantifications. N.G. assisted with experimental design, DART-seq and bioinformatics. X.L. assisted with preparation of primary neurons. R.M. wrote the code for image acquisition and neuronal survival analysis. E.M.T. created all knock-in HEK293T cells and iPSCs, in addition to integrating differentiation cassettes for all iPSC lines. M.B. and E.M.T. maintained iPSCs and differentiated iNeurons for survival experiments. M.B. performed and analyzed iNeuron survival studies. S.J.B. and M.M. assembled figures and wrote the manuscript. S.J.B., M.M., E.M.T., M.B., and N.G. edited the manuscript.

