## Supplemental Information for "RNA methylation influences TDP43 binding and disease pathogenesis in models of amyotrophic lateral sclerosis and frontotemporal dementia"

**Title**

Sami Barmada

University of Michigan

Department of Neurology

109 Zina Pitcher Place, 4019 BSRB

Ann Arbor, MI 48109

### Supplemental methods

#### CFTR minigene assay

HEK293 cells were co-transfected with EGFP or TDP43(WT) and the *CFTR* minigene using Lipofectamine 2000. 48h post-transfection, total RNA was extracted using Trizol according to manufacturer's protocol. cDNA was synthesized using the Bio-Rad iScript cDNA synthesis kit according the manufacturer's protocol, and RT-PCR was accomplished as described previously[1].

#### Purification of recombinant TDP43

TDP43(WT) was expressed in BL21 DE3 E. coli cells from the plasmid pE-6xHis-SUMO-TDP43(WT) (a gift from Dr. James Shorter). Induction was carried out with 1 mM isopropyl-b-D-1-thiogalactopyranoside and cells were grown at 15°C for 16h. Cell pellets were resuspended in lysis buffer (50 mM HEPES, 2% Triton X-100, 300 mM NaCl, 5% glycerol, 50 mM imidazole, 2 mM BME, EDTA-free protease inhibitor cocktail, 5 mM pepstatin, and 20 mg/mL lysozyme) and incubated on ice for 30 minutes. Following sonication on ice, cell lysates were centrifuged for 20 min at 11,000 x g at 4°C. Recombinant protein was purified by binding to Ni-NTA resin (Qiagen #30210), rinsed with 25 mL of wash buffer 4 times (50 mM HEPES, 2% Triton X-100, 300 mM NaCl, 5% glycerol, 50 mM imidazole, and 2 mM BME), and released with 3 mL of elution buffer (50 mM HEPES, 500 mM NaCl, 300 mM imidazole, 5% glycerol, and 5 mM DTT) at room temperature (RT), collecting five 2 mL fractions. Protein was dialyzed twice for 1h in 1 L of final buffer (50 mM HEPES and 500 mM NaCl), and dialyzed once in 1 L of final buffer overnight at 4°C.

#### Electromobility shift assays (EMSAs)

Binding assays were performed with purified full-length recombinant TDP43 protein and either unmodified or m6A modified ssRNA labeled probes tagged with a 5' 800nm infrared (IR) moiety (IDT; **Table S4**). Binding reactions were performed in binding buffer (12.5 mM HEPES, pH 7.8,

50 mM KCl, 2.5 mM MgCl2, 0.5 mM TCEP, 25 mg/mL BSA, 0.01% NP-40) with 50% glycerol, 1 mg/ml poly-dIdC, 500 pM of labeled probe, and recombinant protein (concentrations indicated in figure legends). Reactions were incubated on ice for 5 min followed by 25 min at RT. Electrophoresis of 6% acrylamide gels were performed at 60V. Images were acquired using the LI-COR Odyssey platform.

### Supplemental figure legends

**Supplemental figure 1: HaloTag insertion in *TARDBP* locus does not affect TDP43 function.** (**A**) *CFTR* minigene splicing assay[7,33,75] in which unmodified or HaloTag-TDP43 HEK293T cells were transfected with the CFTR minigene along with EGFP or TDP43-EGFP then analyzed by PCR amplification to measure functional TDP43. Correct TDP43 mediated splicing of the reporter results in exon 9 exclusion. (**B**) Additional dot blots for total RNA (detected by methylene blue) or m6A-modified RNA (detected by anti-m6A antibody) isolated by immunoaffinity purification of endogenous HaloTag-TDP43 or exogenous HaloTag using Synaptic Systems anti-m6a (#202003) or (**C**) Millipore Sigma anti-m6A (ABE572-I-100UG) antibodies.

**Supplemental figure 2: Intronic regions of methylated TDP43 targets are devoid of m6A sites.** Absolute count (**A**) and relative distribution (**B**) of C-T transitions indicative of m6A modifications located within the coding sequence (CDS), 5' untranslated region (UTR), 3' UTR, and introns in cells expressing APOBEC1-YTH and APOBEC1-YTHmut. **(C)** Iterative topological mapping plot depicting the likelihood of finding a TDP43 binding site (as determined by CLIP-seq, Hallegger et al. 2021[2]) within the vicinity of a C-T transition (identified by DART-seq, this study), located at position 0. The magnitude and direction of the slopes for lines corresponding to each gene region indicates a positive relationship between TDP43 binding sites and m6A sites primarily within the 3' UTR, but also within the CDS. (**D**) In the subset of genes exhibiting a TDP43 binding site within 20nt of an m6A site (n=321; grey shading in **C**), this association was most often detected within the 3' UTR (237/321 genes, or 74%). (**E**) The 237 genes showing TDP43 binding sites and m6A sites within the 3'UTR were strongly enriched for genes whose expression is regulated by TDP43, as determined by comparison with the transcription factor loss-of-function GEO database.

**Supplemental figure 3: Methylated TDP43 targets are enriched in RNA binding and homeostasis.** (**A**) Gene ontology for biological processes enriched in TDP43 substrate RNAs (n=1699) common to those identified in this study and Hallegger *et al.* 2021[2]. (B) Protein-protein interaction network analysis[3] for shared TDP43 targets shows enrichment of targets associated with ribosome (**C**), long term potentiation (**D**), VEGF signaling (**E**), and RNA transport (**F**).

**Supplemental figure 4: m6A modifications alter TDP43 binding capabilities**. (**A**) qRT-PCR cycle threshold (Ct) values from HaloTag-TDP43 HEK293T cells expressing mCherry-TBR or mCherry-mTBR. (**B**) Normalized RFP (mCherry) intensity in primary neurons expressing WT (mCherry-TBR) or mutant (mCherry-mTBR) reporters together with EGFP or TDP43-EGFP. mCherry-TBR+GFP n= 160, mCherry-TBR+TDP43(WT)-GFP n= 58, mCherry-mTBR+GFP n= 105, mCherry-mTBR+TDP43(WT)-GFP n= 44. (**C**) Electromobility shift assay (EMSA) demonstrating binding of recombinant TDP43(WT) to 14nt probe modeled after the TDPBR, with and without m6A modification. Probe concentration was kept constant at 500 pM. (**D**) Percent bound m6A modified (red) or unmodified (black) RNA probe. Data in **A** and **B** plotted as mean $\pm$ SD, color coded by biological replicate. ns = not significant; *p < 0.05, **p < 0.01, ***p < 0.001, ****p < 0.0001; one-way ANOVA with Tukey's test. Data in (**C**) plotted as nonlinear regression with Hill slope. ***p < 0.0009; extra-sum-of-squares F test.

**Supplemental figure 5: TDP43-mApple expression is proportional to intensity and toxicity.** (**A**) Correlation of single-cell TDP43 protein levels, determined by immunostaining, and RFP fluorescence in neurons overexpressing TDP43(WT)-mApple. Black: transfected cells, n= 144; grey: non-transfected cells, n= 20. (**B**) Neurons were stratified into 5 quintiles of equal cell number based on TDP43(WT)-mApple intensity, and their risk of death compared via Cox proportional hazards analysis. n= 7016 for mApple + NT; n= 1254 or 1255 cells per quintile. TDP43(WT)-mApple expression is significantly more toxic at all quintiles compared to mApple control: Low HR= 1.38, Med-Low HR= 1.85, Med HR= 2.18, Med-High HR= 2.47, High HR= 2.89. ***$p < 2.0 \times 10^{-16}$ for all quintiles, Cox proportional hazards analysis. (**C**) Density plot of normalized RFP fluorescence illustrating separation of TDP43(WT)-mApple expression quintiles. Inset graphs correspond to neuronal survival in each quintile, same data as displayed in **B**. ***$p < 2.0 \times 10^{-16}$ for all quintiles, Cox proportional hazards analysis.

**Supplemental figure 6: Knockout of m6A pathway components modulates TDP43 toxicity.** (**A**) Knockout of m6A writers Mettl3 (***$p < 2.0 \times 10^{-16}$, HR= 0=2.08; #p= 0.073, HR= 1.06; **$p = 1.45 \times 10^{-3}$, HR= 1.12), Mettl14 (**B**, ***$p < 2.0 \times 10^{-16}$, HR= 2.27; *p= 0.025, HR= 1.08), or Wtap (**C**, ***$p < 2.0 \times 10^{-16}$, HR= 2.08, *p= 0.036, HR= 1.13) does not significantly increase TDP43 dependent toxicity. Knockout of m6A eraser Fto (**D**, ***$p < 2.0 \times 10^{-16}$, HR= 2.0; †$p = 1.31 \times 10^{-8}$, HR= 1.25) does not significantly increase toxicity, but knockout of Alkbh5 (**E,** ***$p < 2.0 \times 10^{-16}$, HR= 2.0; †$p = 3.86 \times 10^{-5}$, HR= 1.23; **$p = 2.87 \times 10^{-3}$, HR= 1.19) significantly increases toxicity. Knockout of m6A reader proteins Ythdf1(**F**, ***$p < 2.0 \times 10^{-16}$, HR= 2.70; †$p = 1.02 \times 10^{-4}$, HR= 1.15; ‡$p = 3.49 \times 10^{-6}$, HR= 1.21 ) and Ythdf3 (**H**, ***$p < 2.0 \times 10^{-16}$, HR= 2.08; **$p = 5.52 \times 10^{-3}$, HR= 1.11; †$p = 5.11 \times 10^{-3}$, HR= 1.14) enhances toxicity, while Ythdf2 knockout (**G**, ***$p < 2.0 \times 10^{-16}$, HR= 2.22; †p= 0.081, HR= 0.95; ‡$p = 1.18 \times 10^{-6}$, HR= 0.84) trends towards neuroprotection. ns= not significant.

Statistical significance determined by Cox proportional hazards, with a minimum of 3 biological replicates per condition.

**Supplemental figure 7: Knockout of additional m6A components at low levels of TDP43 expression is toxic.** (**A**) Knockout of m6A writers Mettl3 (***p= 3.42 x$10^{-6}$, HR= 1.33; #p= 0.056, HR= 1.15); or Mettl14 (**B**, ***p< 2.00 x$10^{-16}$, HR= 1.67;) does not significantly increase toxicity, but Wtap knockout (**C**, ***p= 2.73 x$10^{-6}$, HR= 1.65; *p= 0.026, HR= 1.30) does increase TDP43-dependent toxicity. Knockout of m6A eraser Fto (**D,** ***p= 6.74 x$10^{-8}$, HR= 1.37) does not significantly increase toxicity despite Alkbh5 (**Fig. 5G**) significantly increasing TDP43-mediated toxicity. Knockout of m6A readers Ythdf1 (**E**, ***p<2.0 x$10^{-16}$, HR= 1.76; *p= 0.010, HR= 1.23) and Ythdf3 (**F**, ***p<7.18 x$10^{-8}$, HR= 1.47; *p= 0.026, HR= 1.22) enhance toxicity. ns= not significant. Statistical significance determined by Cox proportional hazards, with a minimum of 3 biological replicates per condition.

**Supplemental figure 8: YTHDF2 immunoreactivity in frontal cortex.** (**A**) Immunostaining of YTHDF2 in control and sALS patient frontal cortex samples. Scale bar= 50µm. (**B**) Quantification of YTHDF2 immunoreactivity in frontal cortex neurons from control (n= 65) and sALS (n= 99) samples. ****p<0.0001, Mann-Whitney test. (**C**) Knockout of YTHDF2 in *C9ORF72* iNeurons reduces toxicity (***p = 1.33x$10^{-6}$, HR= 7.36; †p= 6.74x$10^{-9}$, HR= 5.21; ‡p= 1.16x$10^{-3}$, HR= 6.90; #p= 0.045, HR= 1.9. *p= 0.031, HR= 3.66). Statistical significance determined by Cox proportional hazards.

**Table S1: Post-mortem samples**

| Diagnosis | Age | Sex | Experiment |
|---|---|---|---|
| Control 1 | 66 | M | m6A array |
| Control 2 | 85 | M | m6A array |
| Control 3 | 68 | M | m6A array |
| sALS 1 | 65 | M | m6A array |
| sALS 2 | 64 | M | m6A array |
| sALS 3 | 65 | M | m6A array |
| sALS 4 | 81 | M | m6A array |
| Control 4 | 76 | F | m6A IHC |
| Control 5 | 88 | M | m6A IHC, YTHDF2 IHC |
| Control 6 | 56 | M | m6A IHC, YTHDF2 IHC |
| ALS 1 | 66 | F | m6A IHC |
| ALS 2 | 64 | M | m6A IHC |
| ALS 3 | 68 | M | m6A IHC |
| Control 7 | 48 | M | YTHDF2 IHC |
| ALS 4 | 76 | M | YTHDF2 IHC |
| ALS 4 | 60 | F | YTHDF2 IHC |
| ALS 6 | 99 | M | YTHDF2 IHC |

**Table S2: Plasmids**

| Construct | Reference/<br>Source |
|---|---|
| pGW1-EGFP | 4,5 |
| pGW1-mApple | 4,5 |

| | |
|---|---|
| pGW1-TDP43(WT)-EGFP | 4,5 |
| pGW1-Halo | 6 |
| pGW1-TDP43-TEV-Halo | 7 |
| pCMV-APOBEC1-YTH | 8 |
| pCMV-APOBEC1-YTHmut | 8 |
| pCaggs-mCherry-TBR | 6,7,9 |
| pE-6xHis-SUMO-TDP43(WT) | 7 |
| *CFTR* minigene | 1 |
| pSpCas9(BB)-2A-GFP | Addgene #48138 |
| pcDNA-flag-YTHDF2 | Addgene #52300 |
| pcDNA-flag-METTL3 | Addgene #53739 |
| pcDNA-flag-METTL14 | Addgene #53740 |
| GIPZ Non-silencing Lentiviral shRNA Control | Horizon Discovery #RHS4346 |
| GIPZ Lentiviral Human YTHDF2 shRNA | Horizon Discovery #RHS4430-200182983 |
| pGW1-YTHDF2-Halo | This paper |
| pGW1-YTHDF2-2A-GFP | This paper |

| Construct | Reference/ Source | Amplicon /Insert/ Target | Sequence (5' to 3') |
|---|---|---|---|
| pCaggs-mCherry-mTBR | This paper | TARDBP 3' UTR, SDM A-G | F: CATTATGCACCACCAAGCCTCTGCAC GCGCTCTC |
| | | | R: GCTTTGCAGGAGGGCTTGAAGCAGAG |
| pSpCas9(BB)-2A-GFP + NeuN sgRNA | This paper | NeuN | F: CACCGACCGTCTGGGTCCCAGCGAT |
| | | | R: AAACATCGCTGGGACCCAGACGGTC |
| pSpCas9(BB)-2A-GFP + Mettl3 sgRNA | This paper | Mettl3 | F: CACCGGCTGGGCTTAGGGCCACTAG |
| | | | R: AAACCTAGTGGCCCTAAGCCCAGCC |
| pSpCas9(BB)-2A-GFP + Mettl14 sgRNA | This paper | Mettl14 | F: CACCGGATTCTTCTGGAGCCTCCTC |
| | | | R: AAACGAGGAGGCTCCAGAAGAATCC |
| pSpCas9(BB)-2A-GFP + Wtap sgRNA | This paper | Wtap | F: CACCGGCCGCCAGTCACACAGGCCG |
| | | | R: AAACCGGCCTGTGTGACTGGCGGCC |
| pSpCas9(BB)-2A-GFP + Fto sgRNA | This paper | Fto | F: CACCGGCTGCACAAAGAGGTCCCCG |
| | | | R: AAACCGGGGACCTCTTTGTGCAGCG |
| pSpCas9(BB)-2A-GFP + Alkbh5 sgRNA | This paper | Alkbh5 | F: CACCGGCCTGCCTTGTAGTTGTCCC |
| | | | R: AAACGGGACAACTACAAGGCAGGCC |
| pSpCas9(BB)-2A-GFP + Ythdf1 sgRNA | This paper | Ythdf1 | F: CACCGGCTGTTTTTGGGCAACCTGG |
| | | | R: AAACCCAGGTTGCCCAAAAACAGCC |
| pSpCas9(BB)-2A-GFP + Ythdf2 sgRNA | This paper | Ythdf2 | F: CACCGGCTGTAGTAACTGGGTAAGT |
| | | | R: AAACACTTACCCAGTTACTACAGCC |
| pSpCas9(BB)-2A-GFP + Ythdf3 sgRNA | This paper | Ythdf3 | F: CACCGGCTCTCCCAAGAGAACTAGG |
| | | | R: AAACCCTAGTTCTCTTGGGAGAGCC |
| pSpCas9(BB)-2A-GFP + Atxn2 sgRNA | This paper | Atxn2 | F: CACCGCAGCAGTTCTCGAGGAGGG |
| | | | R: AAACCCCTCCTCGAGAACTGCTGC |

**Table S3: Antibodies**

| Target | Source | Catalog number | Antibody registry number | Species | Dilution |
|---|---|---|---|---|---|
| TDP43 | Proteintech | 10782-2-AP | AB_2892214 | Rabbit | 1:100 |
| NeuN | Cell Signaling Technologies | 24307T | AB_2651140 | Rabbit | 1:500 |
| YTHDF2 | Proteintech | 24744-1-AP | AB_2687435 | Rabbit | 1:100 |
| m6A | Millipore Sigma | ABE572 | AB_2892213 | Rabbit | 1:500 |
| m6A | Cell Signaling Technologies | 56593S | AB_2799515 | Rabbit | 1:500 |
| m6A | Millipore Sigma | ABE572-I-100UG | AB_2892214 | Rabbit | 1:500 |
| HaloTag ligand (JF646 dye) | Promega | GA1120 | N/A | N/A | 1:20,000 |
| Anti-rabbit Alexa Fluor 488 (secondary antibody) | ThermoFisher | A-11034 | N/A | Goat | 1:250 |
| Anti-rabbit HRP (secondary antibody) | Jackson ImmunoResearch Labs Inc. | 111-035-003 | N/A | Goat | 1:10,000 |

**Table S4: Oligonucleotides**

| Target | Source | Sequence (5' to 3') |
|---|---|---|
| mCherry | This paper | F: ATGGTGAGCAAGGGCGAGGA |
| | | R: GATCTCGAACTCGTGGCC |
| *CFTR* | Ayala *et al.*, 2006 | F: caacttcaagctcctaagccactgc |
| | | R: taggatccggtcaccaggaagttggttaaatca |
| Clip34nt_unmodified | IDT | 5IR800CWN-AGAGACUUGGUGGU |

| | | |
|---|---|---|
| Clip34nt_m6A | IDT | 5IR800CWN-AGAG/iN6Me-A/CUUGGUGGU |
| N_TARDBP_F | IDT | CACCGGAAATACCATCGGAAGACGA |
| N_TARDBP_R | IDT | CACCGGGGCTCATCGTTCTCATCTT |

**Table S5: Cell lines**

| Name | Source | Identifier |
|---|---|---|
| HEK293T | ATCC | CRL-3216 |
| Primary rat neurons | University of Michigan Unit for Laboratory Animal Medicine | |
| Human iPSCs | University of Michigan ALS Repository | 1021, 0883, 0312 |
| | Cedars Sinai iPSC Repository | Cs29i, Cs29 |

**Table S6: Human iPSC lines**

| Line | Identifier | Gene | Mutation | Sex | Age at biopsy | Age at onset | Tag | Integrated cassette |
|---|---|---|---|---|---|---|---|---|
| 1021 | WT | *TARDBP* | - | F | 54 | - | TDP43-Dendra 2 (C) | CLYBL:TO-hNgn1/2[6] |
| 1021 | M337V | *TARDBP* | Isogenic M337V[10] | F | 54 | - | TDP43-Dendra 2 (C) | CLYBL:TO-hNgn1/2[6] |
| 312 | C9ORF72 (#1) | *C9ORF72* | HRE | M | 52 | 54 | - | CLYBL:TO-hNgn1/2[6] |
| 883 | C9ORF72 (#2) | *C9ORF72* | HRE | M | 49 | 51 | - | CLYBL:TO-hNgn1/2[6] |
| Cs29i | C9ORF72 (#3) | *C9ORF72* | HRE | M | 47 | NA | - | CLYBL:TO-hNgn1/2[6] |
| Cs29i | C9ORF72 ISO (#3) | *C9ORF72* | Isogenic corrected[11] | M | 47 | NA | - | CLYBL:TO-hNgn1/2[6] |

NA: not assessed; HRE, hexanucleotide repeat expansion mutation; (C), carboxyl terminus

Supplemental Fig. 1 HaloTag insertion in *TARDBP* locus does not affect TDP43 function

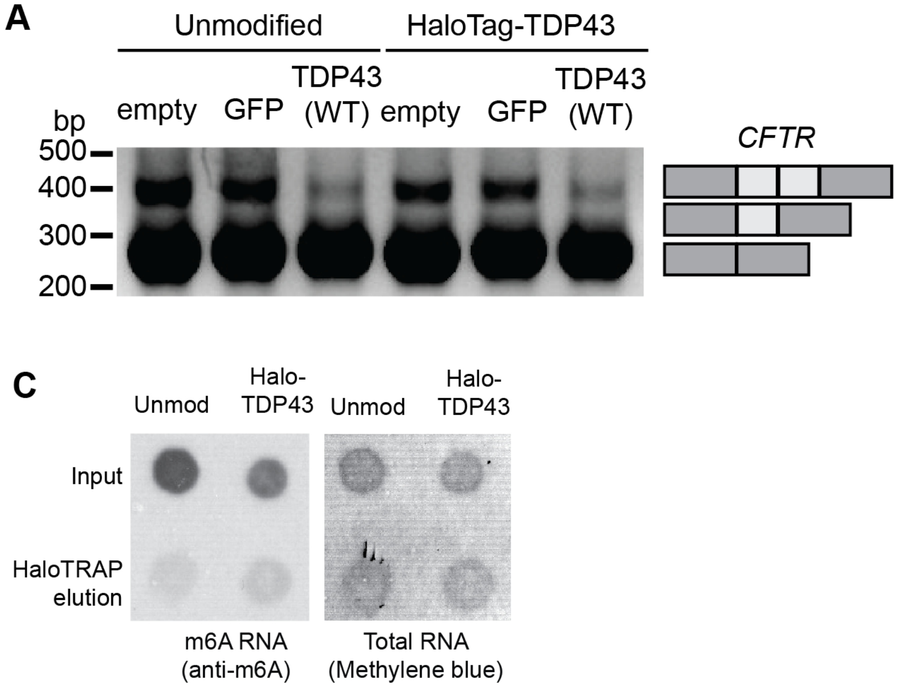

Supplemental Fig. 2: TDP43 binding sites coincide with m6A sites in the 3'UTR

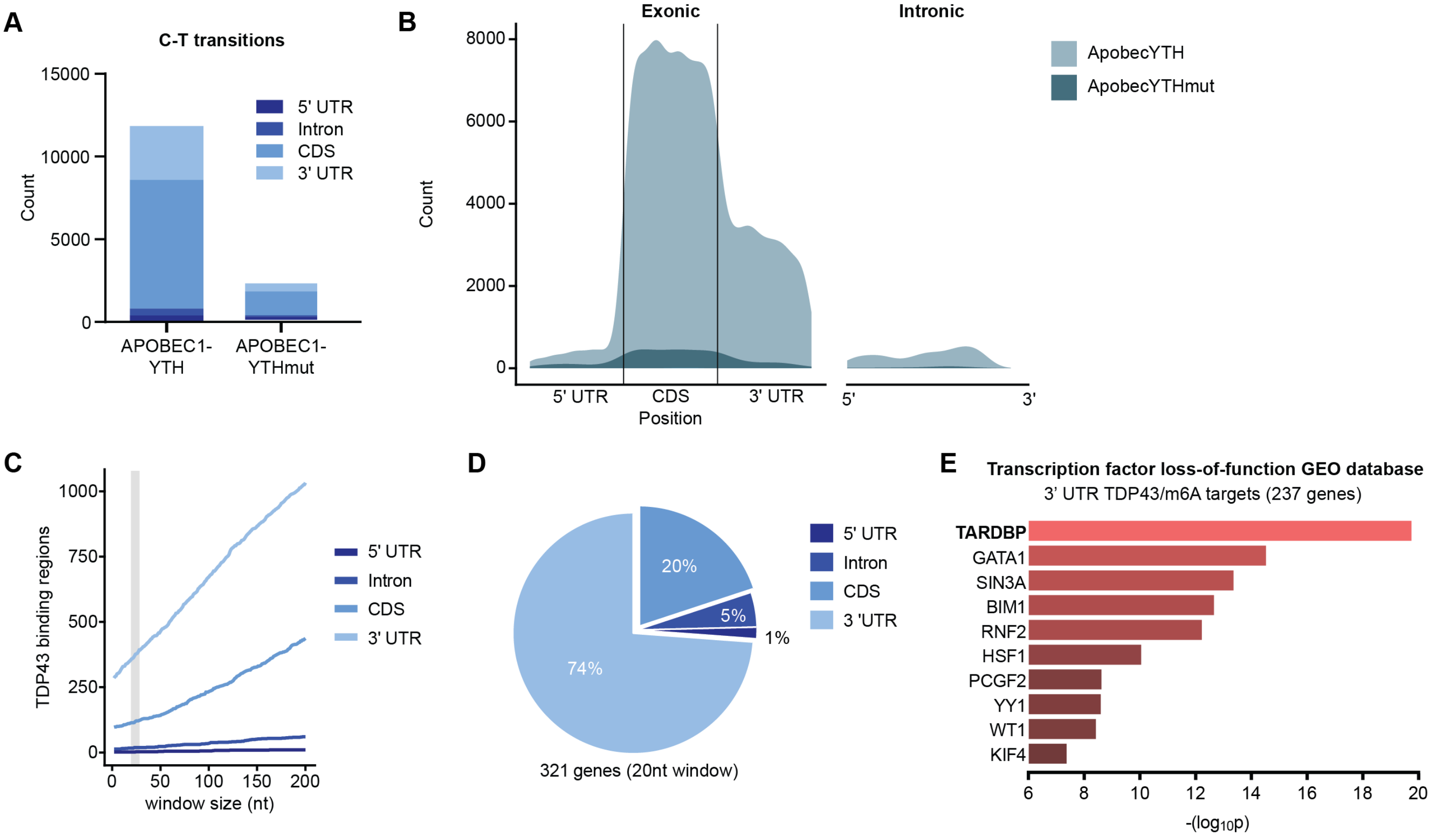

Supplemental Fig. 3: Methylated TDP43 targets are enriched in RNA binding and homeostasis

**A**

**Biological process GO** for methylated Hallegger and DART-seq TDP43 targets (n=1699)

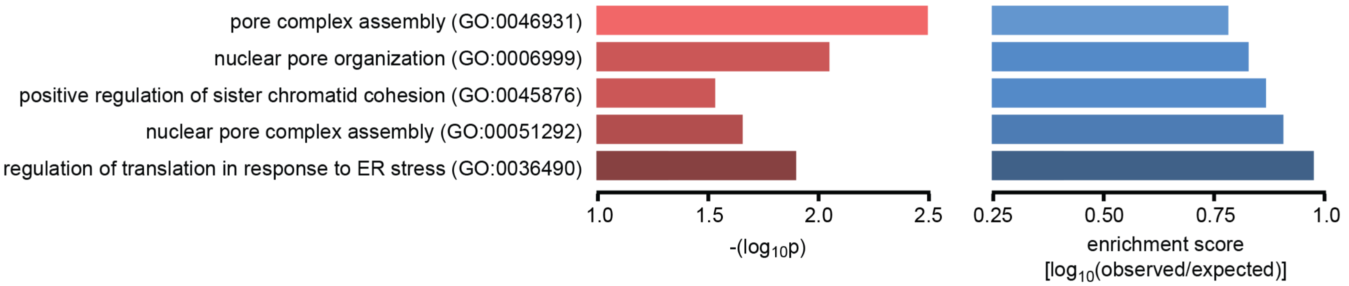


**B** **Protein-protein interaction (PPI) network** analysis for conserved TDP43 targets (n=1699)

| cluster ID | # | examples | top KEGG pathway |
|---|---|---|---|
| 1 | 24 | CDK1, hnRNPC, HSPA4, HSPA5, XPOT | cell cycle |
| 2 | 20 | BCL2, CHEK1, CREB1, TAF15, TP53 | apoptosis |
| 3 | 20 | EIF5, NOP58, RPL11, RPL17, RPS5 | ribosome |
| 4 | 18 | CARM1, CREBBP, ATF4, ETS1, HSF1 | long-term potentiation |
| 5 | 17 | DPHA2, LAMB1, CAPNS1, CDC42 | VEGF signaling |
| 6 | 17 | NUP107, NUP153, NUP214, TNPO1 | RNA transport |
| 7 | 15 | DCTN2, DCTN4, DYNC1H1, CAPZA1 | vasopressin signaling |
| 8 | 15 | ARFGEF1, ADCY6, UBQLN1, CACNB2 | GABAergic synapse |
| 9 | 14 | CDC6, DNA2, POLD2, POLE2, POLG | DNA replication |
| 10 | 14 | CNOT4, PTBP1, EIF4G1, CASP7 | viral infection |

**C**

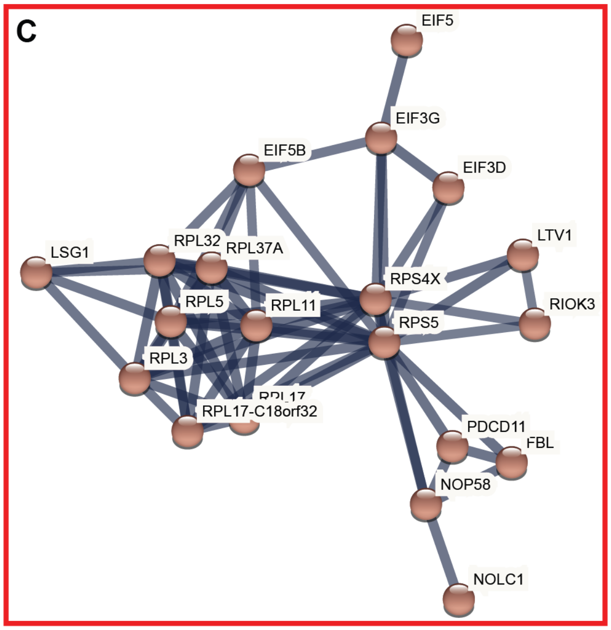


**D**

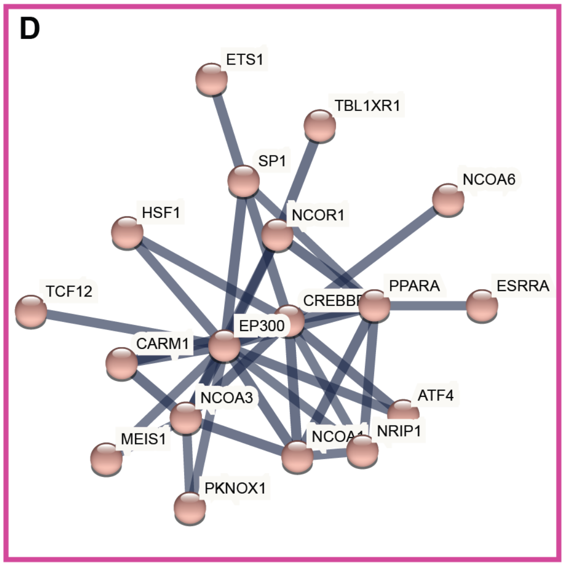


**E**

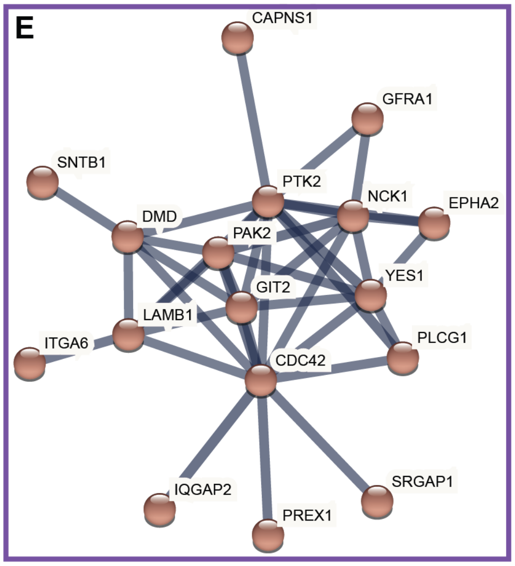


**F**

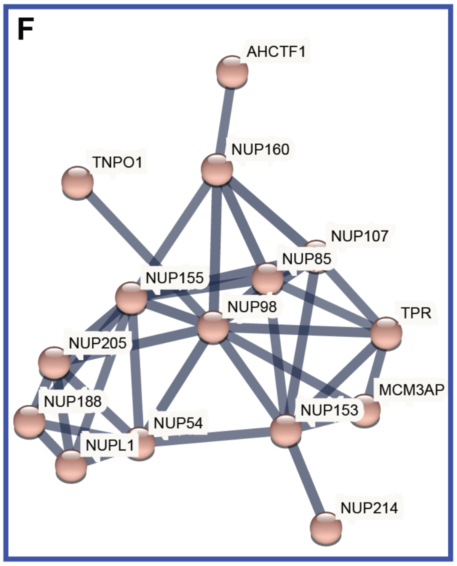

Supplemental Fig. 4: m6A modifications alter TDP43 binding capabilities

A

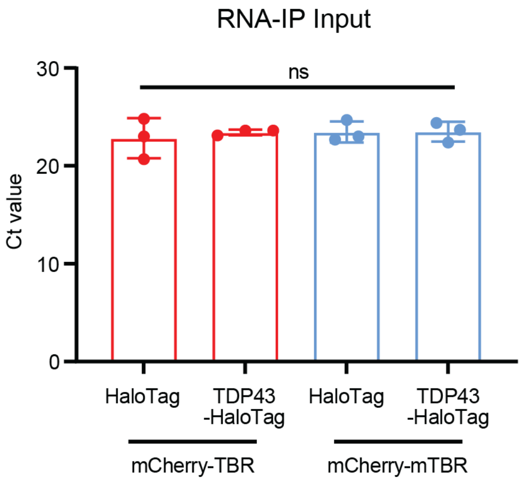


B

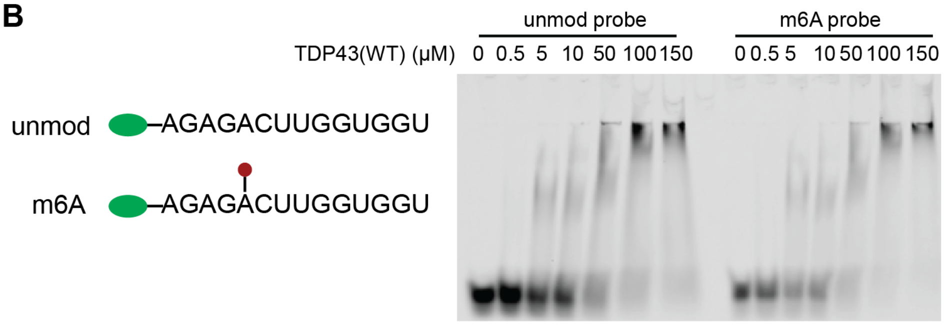


C

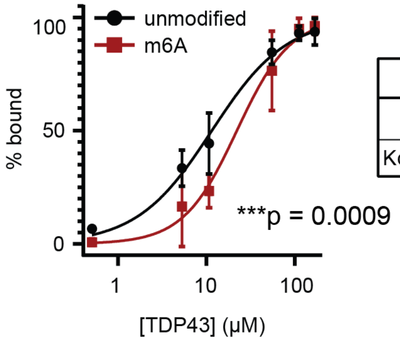


| | unmodified | m6A |
|---|---|---|
| h | 1.041 | 1.453 |
| Kd (μM) | 10.74 | 20.39 |

D

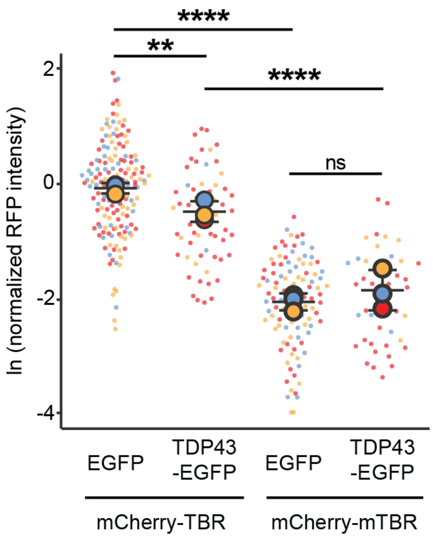

Supplemental Fig. 5: TDP43-mApple expression is proportional to intensity and toxicity

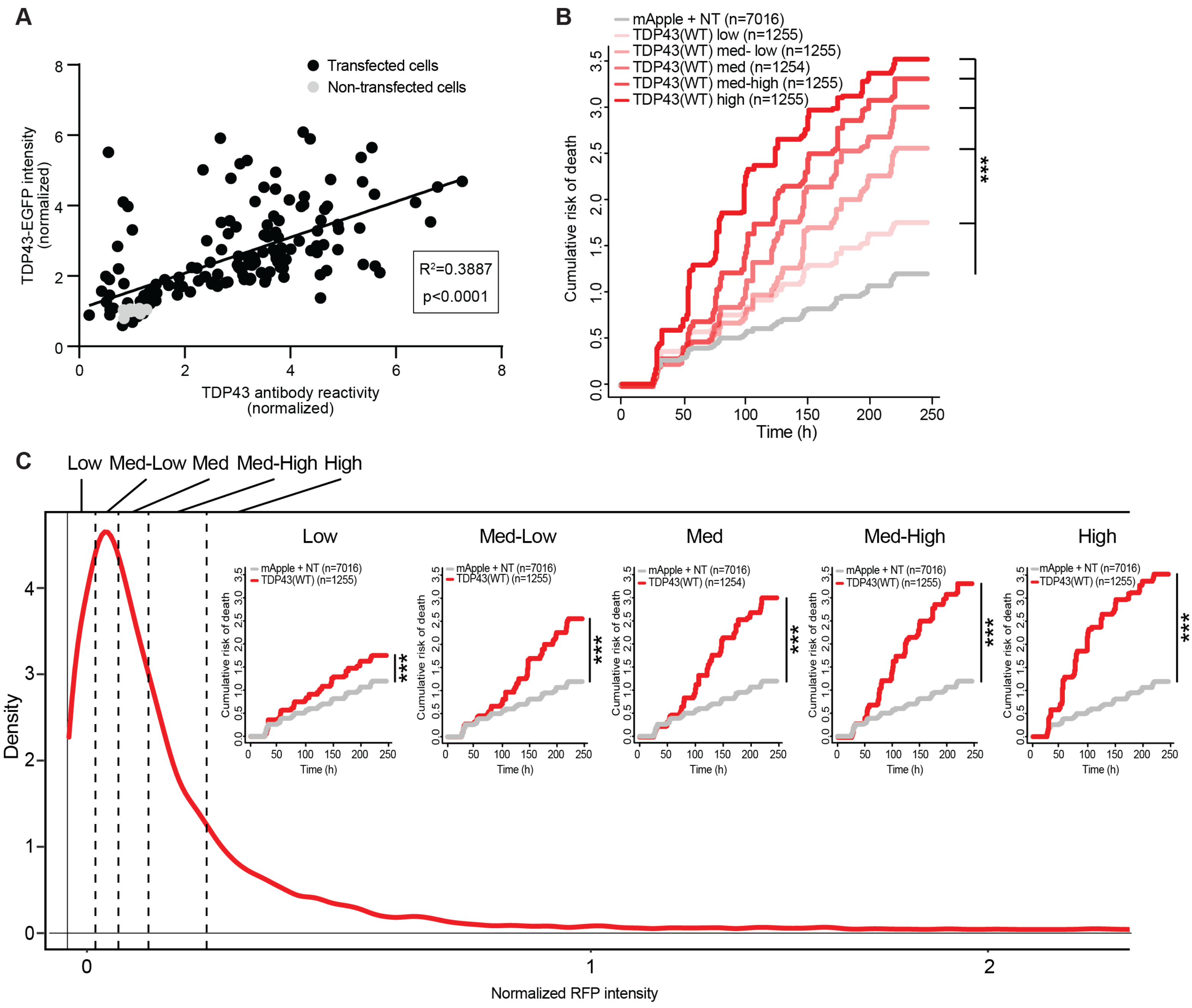

Supplemental Fig. 6: Knockout of m6A pathway components modulates TDP43 associated toxicity

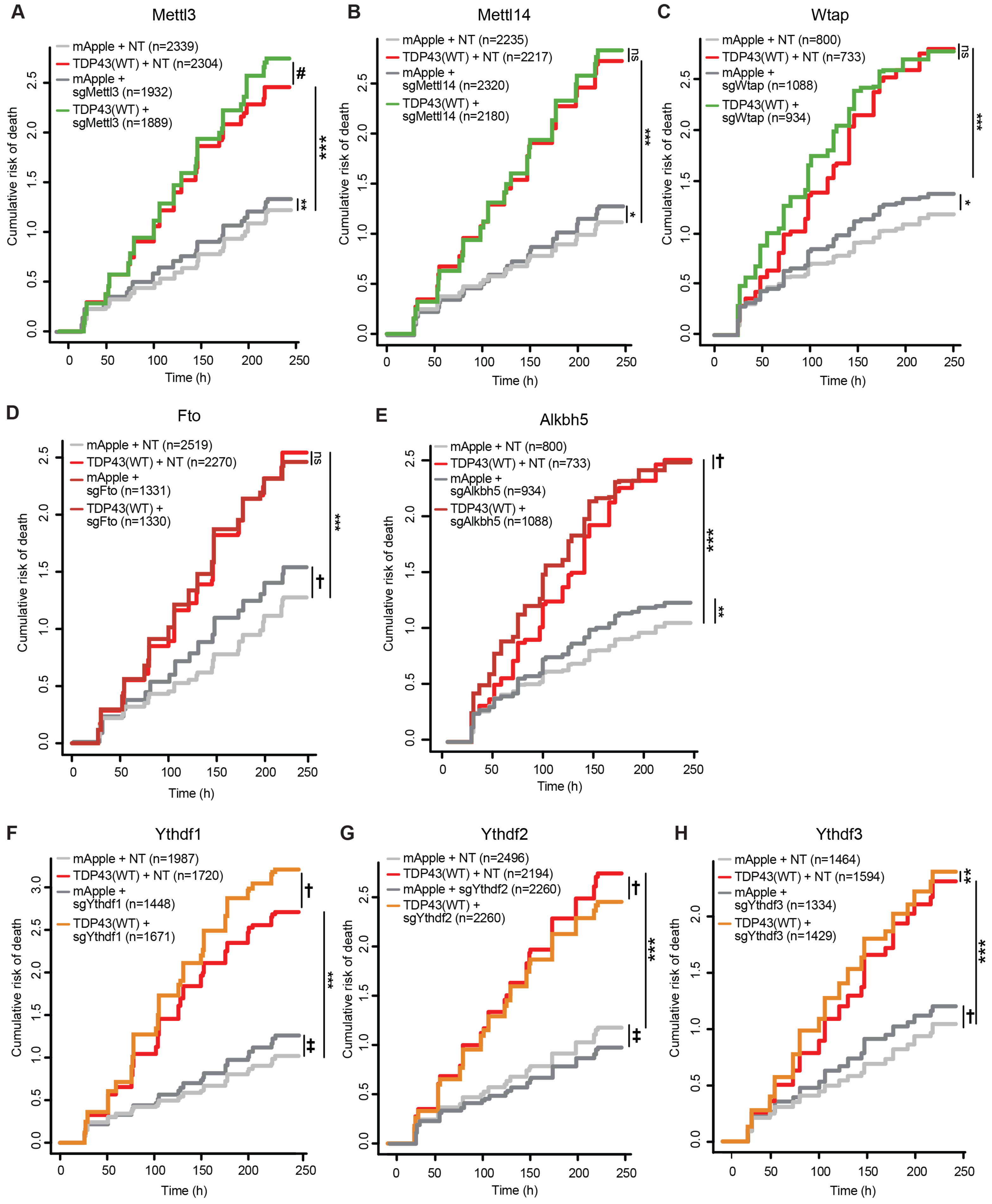

Supplemental Fig. 7: Knockout of additional m6A components at low levels of TDP43 expression is toxic

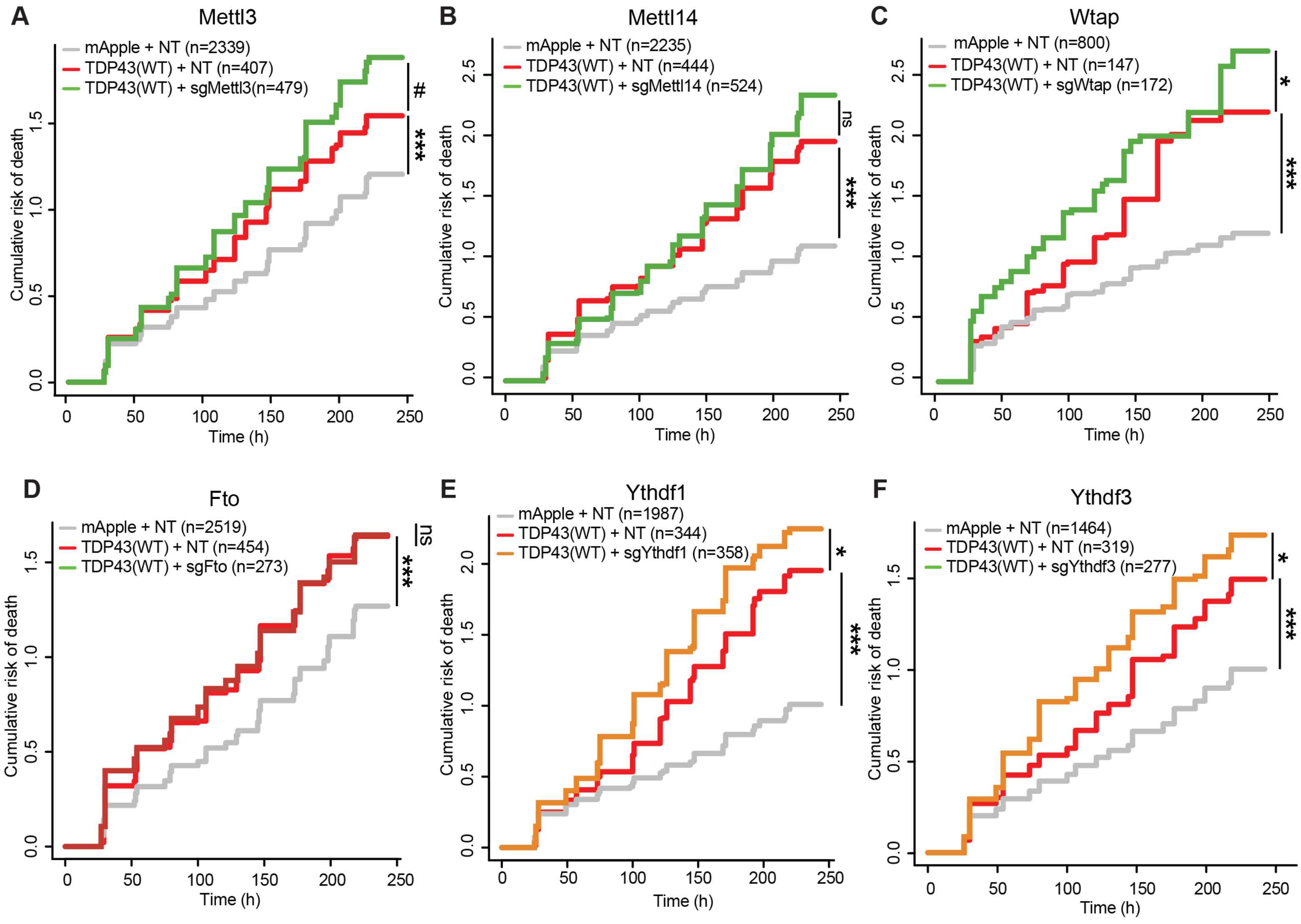

Supplemental Fig. 8: YTHDF2 immunoreactivity in frontal cortex

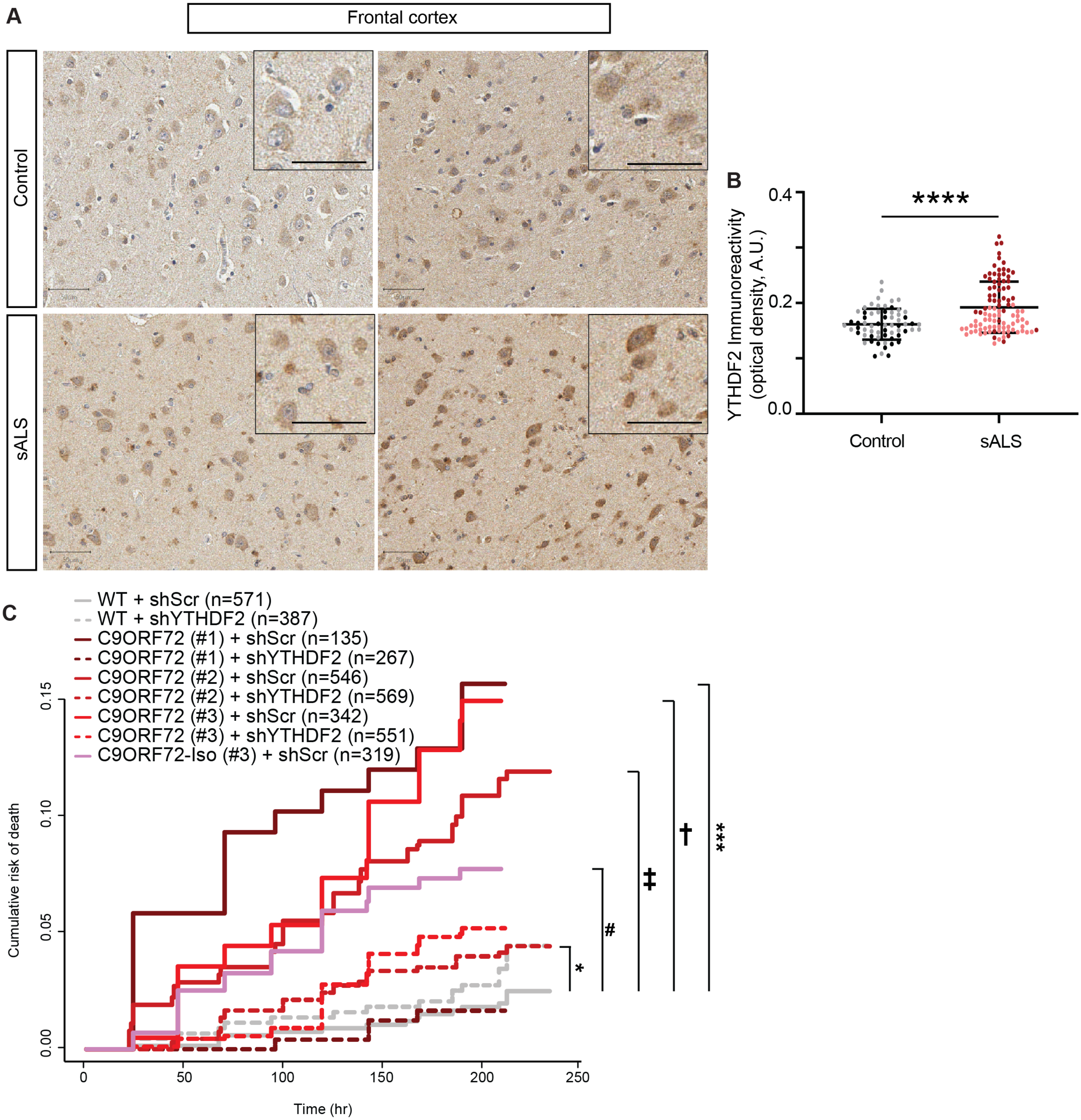
